# Microstimulation of the human dopaminergic midbrain enhances outcome-related happiness and reduces loss seeking

**DOI:** 10.64898/2026.05.15.723537

**Authors:** Mar Yebra, Hristos Courellis, Sophia Cheng, Clayton Mosher, Zhongzheng Fu, Michele Tagliati, Adam Mamelak, Ueli Rutishauser

## Abstract

Midbrain circuits are thought to govern reward-guided behavior and momentary happiness, yet causal evidence in humans remains scarce. Here, we show that focal microstimulation of the substantia nigra pars compacta alters both decision making and momentary happiness in patients undergoing awake deep-brain stimulation surgery for Parkinson’s disease. During a gambling task with interleaved momentary happiness ratings, stimulation delivered at outcome reduced risk-seeking in the loss domain without increasing overall gambling probability. Stimulation improved choice efficiency, guiding behavior toward gambles with higher expected value and thereby increasing cumulative earnings. Computational modeling revealed selective changes in loss processing, with reduced loss weighting and more linear loss utility, while gain processing remained unchanged. In parallel, stimulation did not tonically elevate happiness but selectively amplified outcome-related happiness changes, scaling with decision value and reward prediction errors. These results provide causal evidence that localized perturbation of human midbrain circuits reshapes valuation, choice, and momentary well-being.

## Introduction

A foundational aspect of human behavior is that people weigh equal losses and gains differently, often placing more weight on the former over the latter (Kai-Ineman & Tversky, 1979; Tom, Fox, Trepel, & Poldrack, 2007; Tversky, Kahneman, & uncertainty, 1992). In addition, people also commonly display diminishing sensitivity for gains compared to losses as outcomes become larger (Tversky et al., 1992; Wakker, 2010). What are the neural mechanisms underlying this salient aspect of how we weigh risks and benefits when making decisions? Converging evidence from neuroimaging, neurophysiology, and computational studies implicates a distributed valuation and decision network encompassing the ventral striatum, ventromedial prefrontal cortex (vmPFC), and midbrain (VTA/SNc) in risky decision making (Bartra, McGuire, & Kable, 2013; Chew et al., 2019; Haber & Knutson, 2010; Kable & Glimcher, 2009; Levy, Snell, Nelson, Rustichini, & Glimcher, 2010; O’Doherty, Rutishauser, & Iigaya, 2021; Schultz, 2016b) However, despite strong correlational evidence, this network is highly distributed and the specific causal contributions of individual nodes—and of distinct decision components—remain little understood, particularly in humans (Kable & Glimcher, 2009; Messimeris, Levy, & Le Bouc, 2023).

Deciphering the role of phasic responses of midbrain nuclei during learning has garnered significant attention. This work has revealed a major role for phasic reward-prediction error (RPE) related responses by dopaminergic neurons in the midbrain substantia nigra (SN) (Lak, Stauffer, & Schultz, 2014, 2016; Schultz, 2016a; Schultz, Dayan, & Montague, 1997), including confirming their causal role in reinforcement-driven learning (Stauffer et al., 2016; Steinberg et al., 2013; Tsai et al., 2009). Human studies complement this picture: BOLD responses in the VTA correlate with Reward Prediction Error (RPE) magnitude (D’Ardenne, McClure, Nystrom, & Cohen, 2008), as do fast-scan cyclic voltammetry measures of dopamine (DA) release in the striatum (Kishida et al., 2016). Much less is known about the role of the midbrain in the decision-making process itself beyond learning. It is well recognized that the midbrain dopaminergic system plays a major role in the decision making process itself (Berridge, 2007; Chew et al., 2019; Collins & Frank, 2014; Palmiter, 2008; Salamone & Correa, 2024), extending beyond learning to support motivated and effortful goal-directed behavior. Within this system, the substantia nigra pars compacta (SNc) constitutes a major hub critically implicated in these processes. For example, increasing tonic DA levels increases risk-taking behavior (Burke et al., 2018; Rigoli et al., 2016), an effect that is of paramount clinical relevance in treating Parkinson’s disease (Smittenaar et al., 2012).

Alterations in dopaminergic signaling are a defining feature of Parkinson’s disease (PD), reflecting progressive degeneration of dopamine-producing neurons in the substantia nigra. This degeneration is anatomically heterogeneous, with more severe cell loss in ventral and caudal regions and relatively preserved populations in dorsal areas of the SN, particularly at earlier disease stages (Damier, Hirsch, Agid, & Graybiel, 1999; Fearnley & Lees, 1991). Consistent with this, electrophysiological recordings in PD patients undergoing deep brain stimulation surgery have successfully identified functional dopaminergic neurons in the human substantia nigra (Batten et al., 2024; Imtiaz et al., 2026; Imtiaz et al., 2024; Ramayya, Zaghloul, Weidemann, Baltuch, & Kahana, 2014; Zaghloul et al., 2009), confirming that functional DA neurons remain at this stage of the disease. Recording and stimulating DA neurons in this patient population, as we do here, therefore allows the study of the causal role of DA neurons in behavior.

These alterations in dopaminergic signaling in patients with PD are also frequently accompanied by changes in decision making under risk. However, these changes are not uniform. Risk preference in PD depends critically on dopaminergic state, task structure, and patient subgroup. Unmedicated PD patients have often been described as being more cautious or risk-averse, whereas dopaminergic therapies—particularly DA agonists—can shift behavior toward increased risky choice and are associated with impulse-control disorders such as pathological gambling (Claassen et al., 2011; Heiden, Heinz, & Romanczuk‐Seiferth, 2017; Rogers, 2011; Timmer, Sescousse, Esselink, Piray, & Cools, 2018). At the same time, the effects of dopamine on loss processing are complex: medicated PD patients have been reported to show reduced loss aversion in effort-based decision-making tasks (Chen, Voets, Jenkinson, & Galea, 2020), whereas other studies have observed enhanced sensitivity to potential losses depending on task demands (Nobis et al., 2023). Together, these findings indicate that dopaminergic perturbations can reshape how prospective gains and losses are weighted, but the direction and computational nature of these changes remain unresolved.

Consistent with a DA-sensitive contribution to loss processing, medicated PD patients show reduced loss aversion relative to controls in effort-based decision-making paradigms (Chen et al., 2020). In parallel, impulse-control disorders—including pathological gambling—are over-represented in PD, particularly in patients receiving dopaminergic therapies, and occur at substantially higher rates than in the general population (Heiden et al., 2017; Santangelo, Barone, Trojano, Vitale, & disorders, 2013). These clinical observations underscore the importance of understanding how dopaminergic perturbations influence valuation and choice.

Beyond influencing decision making, dopamine-dependent valuation processes also bear on another longstanding question: how do objective outcomes relate to subjective well-being? Philosophical and economic accounts have long emphasized that happiness is not reducible to material payoffs, and empirical work shows only a shallow, nonlinear relationship between income and experienced happiness, with life evaluation and emotional well-being dissociating at higher incomes (D. Kahneman & Deaton, 2010), whereas more recent large-scale analyses report continued increases without a clear upper bound (Killingsworth, 2021). These observations underscore that the mapping from objective value to subjective experience is not fixed, but depends on measurement, context, and underlying mechanisms, motivating a mechanistic test of how causal modulation of midbrain circuits shapes both decision quality and momentary happiness. Equally importantly, there is also extensive literature linking midbrain activity, and in some instances specifically DA levels, to subjective momentary happiness (Rutledge, Skandali, Dayan, & Dolan, 2015; Rutledge, Skandali, Dayan, & Dolan, 2014). This work shows that the best predictor of momentary happiness is RPEs (Rutledge et al., 2014). While the relationship between midbrain activity and/or DA levels and risky decision making as well as momentary happiness is well established, little is known about the causal role of the midbrain in this process.

Here, we employed SNc microstimulation to examine the casual role of SNc activity in both risky decision-making and the assessment of subjective momentary well-being. We did so using intra-operative microelectrode recording in conjunction with behavioral testing in 24 patients with PD undergoing implantation of a Deep Brain Stimulation (DBS) device in the subthalamic nucleus (STN) to treat motor symptoms of their disease. Subjects performed a well validated economic decision-making task that includes a happiness rating component (Rutledge et al., 2015). We sought to test three hypotheses. First, given the important role of SNc activity in decision making, we hypothesized SNc stimulation would modify choice behavior. Second, because human behavior in this task is well described by prospect theory, we hypothesized that we could quantify the resulting stimulation-related changes in behavior by changes in specific parameters in this model. Third, we proposed that stimulation could modulate moment to moment perceived subjective happiness.

## Results

### Task and Subjects

Subjects performed a probabilistic risky decision-making task. In each trial, subjects were offered a choice between a certain option (without gambling) or a gamble option with two possible outcomes that were equally likely (Figure 1a; the numbers shown on the screen are multiplied by 100 to make them more salient). After choosing either the certain or the gambling option, subjects were shown the amount of reward they received (outcome period’), which was either the amount offered for sure or one of the two equiprobable gamble outcomes, depending on the decision the subject made earlier in the trial. In addition, every 3 trials, subjects were asked “How happy are you right now?”, which they answered by moving a slider using a track ball mouse (Figure 1a, b). Happiness responses were converted to a −100 to 100 scale by linearly rescaling each cursor position on the rating line, such that the left endpoint (very unhappy) = −100, the midpoint = 0, and the right endpoint (very happy) = 100. Subjects were instructed that their goal is to accumulate as many total reward points as possible.

**Figure 1.**
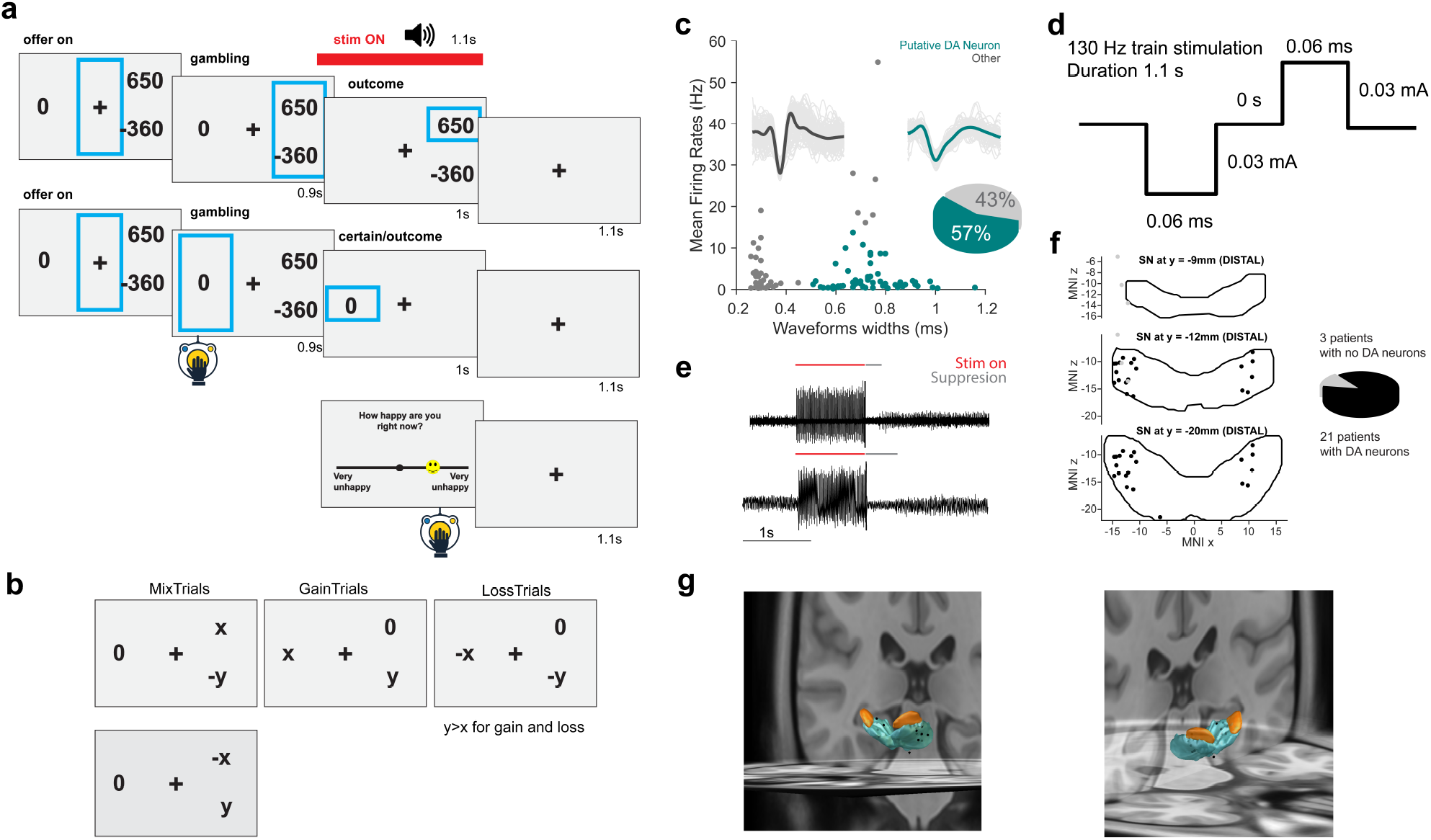
Task, Neural Recordings, and Stimulation Parameters. **(a)** Schematic of the task. In each trial, patients chose between a certain reward and a monetary gamble. After a 1-second delay, the outcome achieved was shown. Every three trials, patients rated their current happiness using button presses. **(b)** Illustration of the three possible types of gamble trials: gain (possible gain vs. 0), mix (gain or loss), and loss (certain loss vs. larger potential loss). **(c)** Neuronal classification based on waveform and baseline firing rate. Putative dopamine (DA) neurons (dark green) exhibited long-duration waveforms and low firing rates; example waveforms are shown for a DA neuron (green) and a fast-spiking GABAergic neuron (dark gray). Light gray lines indicate waveforms of all detected units for each cell (n=101). **(d)** Stimulation parameters and waveform. Stimulation consisted of biphasic pulses delivered in short trains. **(e)** Sample voltage trace showing band-pass filtered signal, highlighting the silent period that follows the stimulation train (red indicates stimulation is ON). **(f)** Electrode locations across subjects in 2D using the Distal atlas showing z and x coordinates in MNI space at the different y levels. **(g)** 3D visualization of electrode locations using Distal atlas Lead-DBS and the (Yu Zhang, Larcher, Misic, & Dagher, 2017) STN atlas, with two viewing angles.

Subjects were 24 patients with PD who underwent implantation of a DBS device in the STN to treat motor aspects of their disease. The experiments reported here took place during the surgery, as part of which patients were awake and underwent microelectrode recordings. During this procedure, a microelectrode is advanced into the substantia nigra (SNc) (Figure 1f-g) to identify the inferior border of the subthalamic nucleus (STN) for clinical purposes (see Methods). We confirmed that the electrode tip was in the SN through microstimulation (Figure 1e, see methods). While subjects performed the task, we applied microstimulation (0.03mA, 130Hz, Figure 1d) through the microelectrode located in the SN. Stimulation was applied during the outcome period for a 1.1s long period of time for a random subset of half of the trials in blocks of 3. We refer to trials that immediately follow an outcome period with stimulation or without stimulation as ‘ON’ and ‘OFF’ trials, respectively.

Analysis of neuronal activity waveform determined whether the tip of the stimulation electrode was near likely dopaminergic neurons. To analyze the recorded neuronal activity, we removed the stimulation artifact from the recorded signal (see Methods, Supplementary Figure 3) and sorted the data to identify putative single units (see Methods). We recorded 101 SN single neurons in total. For each recorded neuron, we determined whether a neuron is likely to be dopaminergic (DA) based on properties of its action potential waveform shape and its mean firing rate (Figure 1c; see methods) (Kamiński et al., 2018a). To increase the likelihood that DA neurons were stimulated in our data, we included in our behavioral analysis only the n=21 sessions in which at least one putative DA neuron was identified (Figure 1c,f; electrode location of patients with DA neurons present are marked with black dots) (see Methods).

### Choice behavior

We first examined whether subjects made risky decisions by weighing the risks and benefits of the offered gambles against each other. To do so, we quantified choice behavior using the difference between the expected value of the gamble (EV; mean of the two possible outcomes) and the value of the certain option (CR). This quantity corresponds to the relative (decision) value of the gamble compared to the certain option (ΔEV), a standard measure in value-based decision-making models. Positive values indicate that the gamble has higher expected value than the certain option, whereas negative values indicate that the certain option is preferable. We found that subjects understood the task: larger EV-CR, correlated with higher probability of subjects gambling (Figure 2a; pooled across stimulation ON and OFF trials; magnitude of EV/CR is divided by 100 relative to what is shown on the screen in Figure 1a). To statistically assess the robustness of this relationship, we used a per-participant aggregated binomial GLM of gamble probability as a function of EV–CR. The mean slope was significantly larger than zero (one-sample *t-*test: *t*_19_ =2.9865; p = 0.008). This analysis shows that choices were influenced by the value of the gamble, as intended.

**Figure 2.**
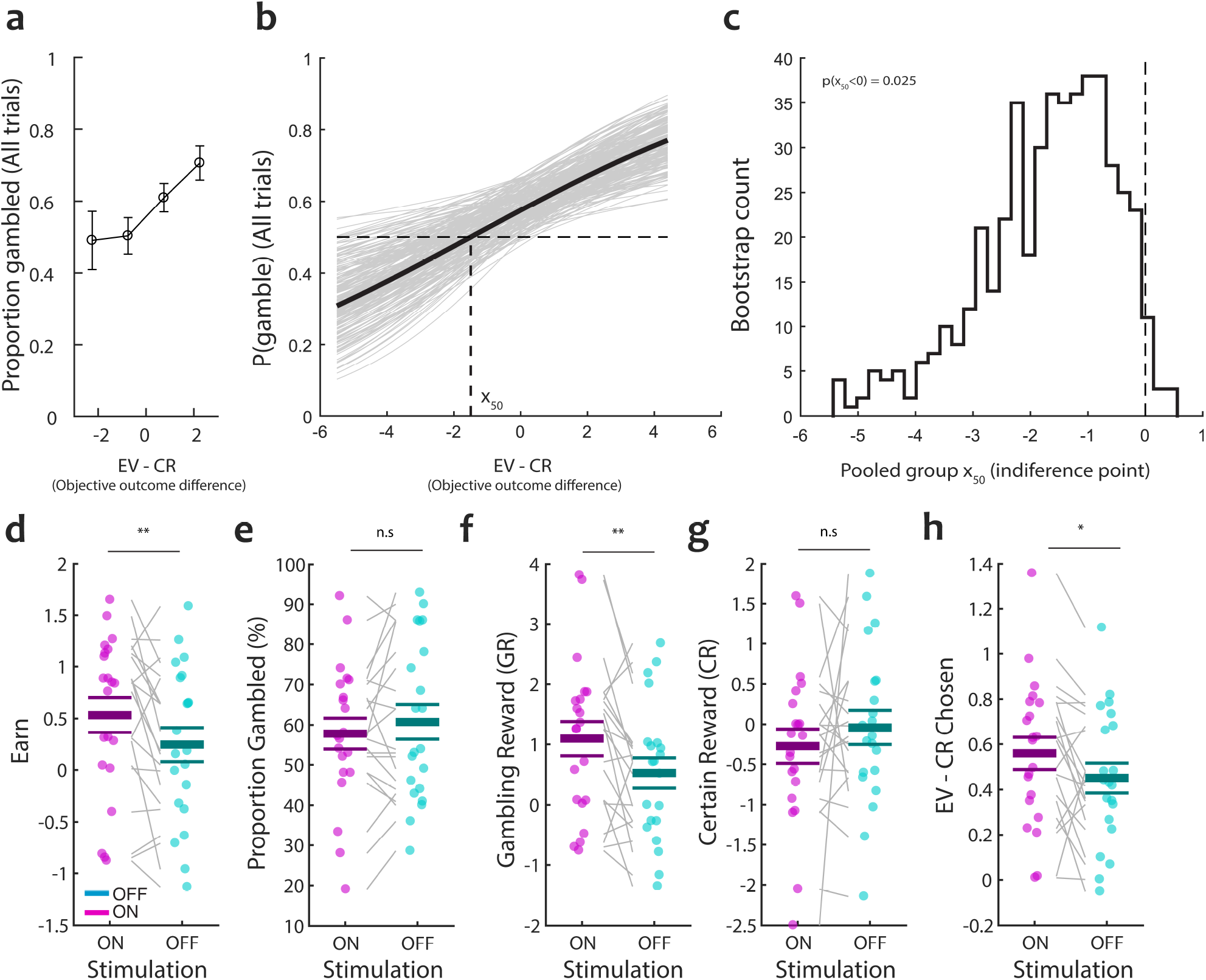
Patient’s are risk-seeking and SNc stimulation increases choice value and earnings. (a) Probability of choosing the gamble as a function of objective value difference (EV−CR), showing group mean ± s.e.m. (b) Psychometric function collapsing across stimulation ON and OFF conditions. Gray lines indicate bootstrap resamples; black line indicates the group-average fit from a logistic model with subject intercepts. The vertical dashed line indicates the indifference point (x_50_). A negative x_50_ reflects loss-seeking behavior in the loss domain. (c) Bootstrap distribution of the pooled indifference point (x_50_). The dashed line indicates x_50_ = 0. The distribution is shifted below zero, consistent with risk-seeking behavior (one-sided bootstrap p = 0.025). (d–h) Participant-level summaries. Each dot represents one participant; gray lines connect stimulation OFF and ON conditions; bars show mean ± s.e.m. (d) Earnings per trial. (e) Overall gambling rate. (f) Gambling reward (GR) on chosen gamble trials. (g) Certain reward (CR) when the gamble was not chosen. (h) EV−CR of the chosen option (choice optimality). Asterisks indicate paired two-tailed tests (*p < 0.05, **p < 0.01; n.s., not significant). Colors: stimulation OFF (cyan), stimulation ON (magenta).

To further characterize choice behavior, we estimated the psychometric function relating gamble probability to the objective value difference between the gamble and the certain option (EV−CR; see methods), pooling across ON and OFF stimulation trials (Figure 2b). The resulting curve (Figure 2b) revealed a systematic bias toward gambling even when the expected value of the gamble was lower than that of the certain option. This was quantified by the indifference point (x_50_), defined as the value of EV−CR at which participants were equally likely to gamble or choose the certain option (‘point of subjective equality’). On average, x_50_ was -1.545 and thus negative, indicating that participants were willing to accept gambles with (small) negative expected value relative to the certain alternative. This type of behavior is consistent with risk-seeking behavior. To assess the robustness of this effect, we performed a bootstrap analysis over subjects and trials (Tibshirani, Efron, & probability, 1993). The distribution of bootstrap estimates of x_50_ was predominantly negative (Figure 2c), with a one-sided bootstrap test confirming that x_50_ was significantly below zero (p = 0.025). Together, these results indicate that altogether, participants exhibited loss-seeking behavior.

We next examined the effect of stimulation on the behavior of the subjects. Participants earned more per trial when stimulation was ON (Figure 2d; paired t-test: *t*_20_ = 2.69; p = 0.01). This effect was the case despite stimulation not affecting the overall probability of subjects to gamble: the average rate of gambling was not significantly different between the ON and OFF conditions (Figure 2e; paired t-test: *t*_20_ = -1.27; p = 0.22). This raises the question of what aspect of the subject’s choice behavior was affected by stimulation. To examine this question, we next examined separately the two different sources from which subjects could gain rewards. If a subject chose not to gamble, the reward they received was referred to as the certain reward (CR), whereas the earnings in trials in which they choose to gamble are referred to as gambling rewards (GR). Decomposing earnings into whether they are due to CR or GR rewards (Figure 2f-g) revealed that subjects received significantly more reward when they choose to gamble during stimulation ON trials (GR; Figure 2f; paired t-test: *t*_20_ = 2.81; p = 0.01). In contrast, when subjects chose not to gamble the reward they received did not differ significantly between stimulation ON and OFF trials (CR; Figure 2g; paired t-test: *t*_20_ = -0.86; p = 0.40). Consistent with this pattern, the EV–CR difference of accepted gambles was larger in stimulation ON than stimulation OFF trials (Figure 2h; paired t-test on subject means: t_20_ = 2.17; p = 0.04). Thus, stimulation did not indiscriminately increase risk taking; rather, participants were more likely to accept gambles when they offered a greater objective advantage over the certain option. This pattern is consistent with increased sensitivity to value differences and more selective, value-guided choice, resulting in higher realized earnings. Importantly, when collapsing across stimulation conditions, participants exhibited a baseline tendency toward risk-seeking behavior in the loss domain, as reflected by a negative indifference point (x_50_ < 0). This indicates that participants were willing to accept disadvantageous gambles relative to the certain option. Against this baseline bias, stimulation improved the quality of choices by increasing sensitivity to objective value differences. Further below, we use prospect theory to examine what aspects of valuation were altered by stimulation to produce these changes in behavior.

### Happiness behavior

We next turned to examining the subject’s happiness ratings, which subjects were asked to provide every 3 trials. We first examined whether the happiness ratings were meaningful. To do so, we fit a regression model (see methods) to examine whether happiness ratings are systematically related to the task variables of the trials that preceded a given happiness rating. We examined received reward, EV, and RPE. The classic model of momentary happiness. shows that these variables predict happiness ratings with high accuracy (Rutledge et al., 2015; Rutledge et al., 2014), making this an appropriate test of the to validate of the ratings provided by our subjects.

Across all trials, happiness ratings were well explained by this model: they explained substantial variance in happiness ratings (R^2^ = 0.30 ± 0.20, mean ± SD). Model fit was quantified within each participant as the Pearson correlation r between observed ratings and fitted ratings across rating time points, and group-level inference was performed by applying Fisher’s r-to-z transform z=tang^-1^(r) and testing whether mean z differed from zero with a one-sample t-test. Model fits were reliably positive across participants (median fitted–observed correlation r = 0.52, interquartile range (IQR) [0.36, 0.63]; Fisher-z group test: t_20_=9.75, p=4.81 10^−9^, mean r from mean z = 0.55). This was also true when examining happiness ratings separately during stimulation ON and OFF trials (OFF-only: mean r=0.77, Fisher-z test: t_18_=6.69, p=2.85 10^−6^; stimulation ON-only: mean r=0.76, t_18_=12.31, p=3.36 10^−10^). Together, these results show that happiness ratings validly reflected moment-to-moment fluctuations in perceived mood.

We next examined the effect of stimulation on the happiness ratings. Figure 3 a-b shows the time course of happiness for an example participant in a single session (straight line). While happiness ratings fluctuate, their average level stays relatively stable within a session. Across participants, mean happiness was comparable between stimulation ON and OFF trials; a paired t-test on the subject z-scored means confirmed no significant difference (ON vs OFF: *t*_20_ = 0.58; p = 0.57). Thus, microstimulation did not produce an overall tonic shift in self-reported effect during the task (Figure 3c).

**Figure 3.**
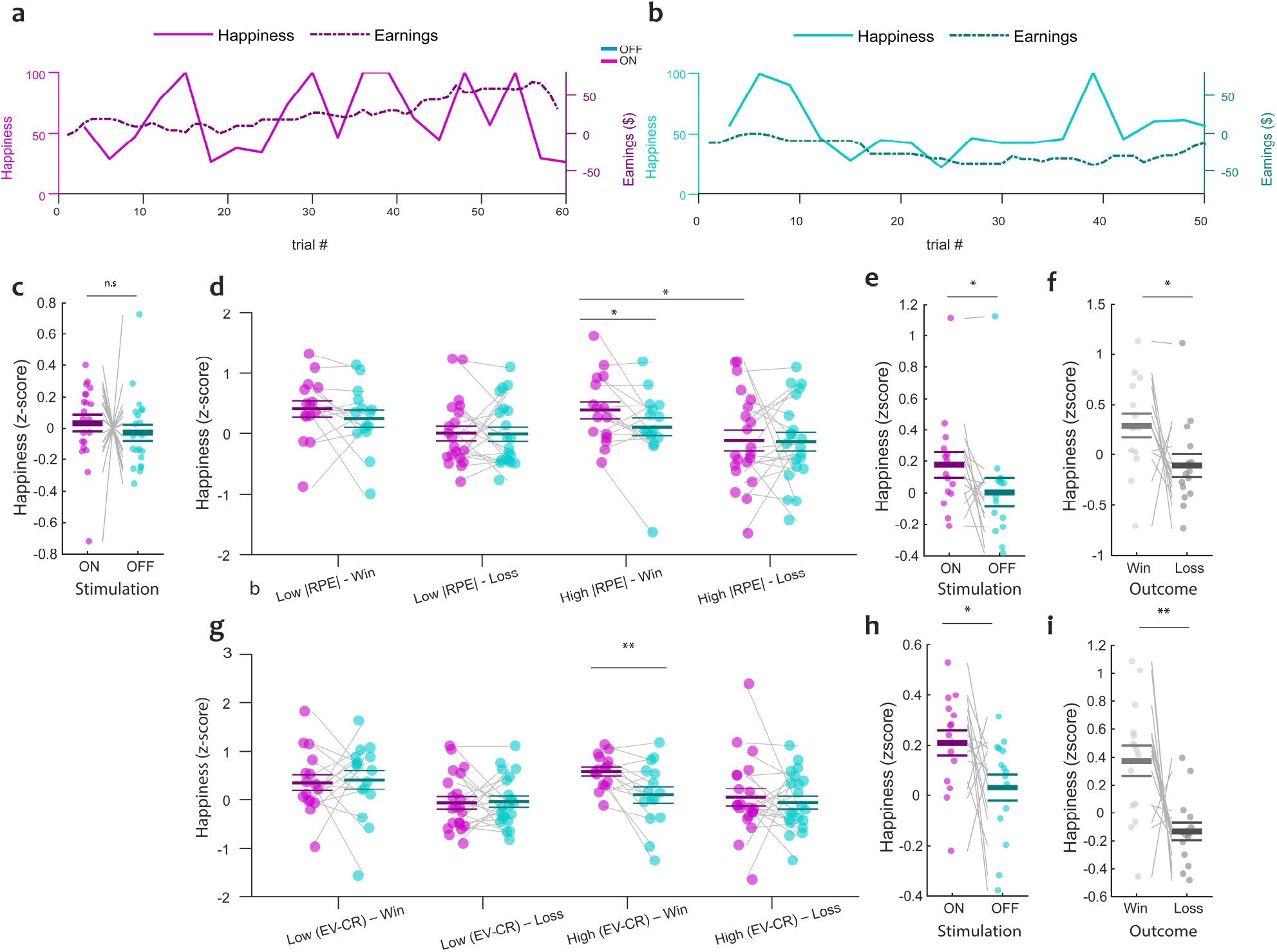
SNc stimulation modulates happiness in relation to prediction error and decision value. (a,b) Example single-session time courses of z-scored happiness (solid line, left axis) and cumulative earnings (dashed line, right axis) during stimulation OFF (a) and ON (b) trials. (x-axis shows within-condition trial order; OFF and ON trials were interleaved in the task). (c) Mean z-scored happiness ratings under stimulation ON versus OFF. (d) (c) Mean happiness as a function of absolute reward prediction error (|RPE|; low/high) and outcome (win/loss) for gamble trials, showing increased happiness under stimulation and after wins. (e,f) Collapsed main effects for the |RPE| analysis. (e) Stimulation increases overall happiness (ON vs OFF). (f) Happiness is higher for wins than losses. (g) Mean happiness as a function of decision value (EV−CR; low/high) and outcome (win/loss), revealing a selective enhancement of happiness for high-value wins under stimulation. (h,i) Collapsed main effects for the EV−CR analysis. (h) Stimulation increases overall happiness. (i) Happiness is higher for wins than losses. Data points represent individual participants; lines connect paired observations. Horizontal bars indicate mean ± s.e.m. Asterisks denote paired two-tailed t-tests (*p < 0.05, **p < 0.01; n.s., not significant). Colors: stimulation OFF (cyan), stimulation ON (magenta).

We next asked whether happiness after a gamble outcome depended on RPE magnitude. Restricting the analysis to happiness ratings that immediately followed trials in which subjects chose to gamble (gamble trials), we classified the RPE associated with the trial preceding each happiness rating as low or high based on a within-subject median split of absolute RPE (|RPE|). A repeated-measures ANOVA on z-scored happiness ratings with within-subject factors stimulation (ON/OFF), RPE magnitude (low/high), and Outcome (win/loss) revealed (Figure 3) significant main effects of stimulation (ON > OFF; F(1,14) = 5.94, p = 0.028) and Outcome (wins > losses; F(1,14) = 5.503, p = 0.0342), but no significant main effect of RPE magnitude (F(1,14) = 2.548, p = 0.1328) nor significant interactions (all p > 0.4). Consistent with the ANOVA main effect of stimulation, happiness ratings were higher during stimulation ON than OFF when pooling across outcome and RPE conditions (Figure 3e; ON: 0.177 ± 0.082; OFF: 0.004 ± 0.091; paired t-test: t_14_ = 2.438, p = 0.0287). Likewise, consistent with the ANOVA main effect of outcome, happiness ratings were higher following wins than losses when pooling across stimulation and RPE conditions (Figure 3f; wins: 0.290 ± 0.119; losses: –0.109 ± 0.113; paired t-test: t_14_ = 2.346, p = 0.0342). When comparing ON versus OFF separately within each RPE × Outcome condition, the stimulation effect was most pronounced for high-|RPE| wins (ON: 0.414 ± 0.149; OFF: 0.088 ± 0.158; t_14_ = 2.385, p = 0.0318), whereas the corresponding contrasts were not significant for low-| RPE| wins, low-|RPE| losses, or high-|RPE| losses (Figure 3d). Similarly, when examining win– loss contrasts separately within each stimulation × RPE cell, the effect was most evident for ON high-RPE trials (win: 0.414 ± 0.149; loss: –0.174 ± 0.213; t_14_ = 2.170, p = 0.0477), with similar but non-significant trends in the remaining cells. Together, these results indicate that, for gamble trials, happiness tracks outcome valence and is elevated by SNc stimulation, with the strongest condition-wise stimulation effect observed following high-RPE magnitude wins.

We next conditioned happiness on the quality of the preceding decision rather than on gambling choice alone. For each happiness rating, we assessed the EV − CR of the trial that preceded the happiness rating. We then median-split trials within subject into those with low- and high-value offers (EV-CR low vs. high; regardless of whether the subjects chose to gamble or not). As expected, happiness ratings were higher following wins than losses (Figure 3g). This pattern held for both the high and low value gamble trials. A repeated-measures ANOVA on z-scored happiness with within-subject factors stimulation (ON/OFF), Value (low/high EV–CR), and Outcome (win/loss) revealed a robust main effect of Outcome (F(1,14) = 9.173, p = 0.009). This win–loss difference did not significantly depend on value (Value × Outcome: F(1,14) = 0.30, p = 0.59), and was confirmed by a paired comparison pooling across conditions (Figure 3i; wins: 0.37 ± 0.11; losses: –0.13 ± 0.06; t_14_ = 3.03, p = 0.009). In addition, we observed a significant main effect of stimulation, with higher happiness ratings during stimulation ON compared to OFF (F(1,14) = 4.74, p = 0.047), indicating that SNc stimulation produces an overall increase in momentary happiness (Figure 3h; ON: 0.210 ± 0.049; OFF: 0.032 ± 0.052; t_14_=2.177, p=0.047). To further examine how this effect manifests across decision contexts, we compared happiness ratings between ON and OFF trials separately for high- and low-value (EV − CR) conditions. While stimulation effects were not significant in most individual condition combinations, the largest increase in happiness was observed following high-value wins (ON: 0.58, OFF: 0.10; t_14_ = 3.05, p = 0.009; Figure 3g). Together, these results indicate that happiness reliably tracks outcome valence, while SNc stimulation produces a general increase in happiness, with the strongest effects observed following high-value wins, consistent with a general modulation of momentary happiness by SNc stimulation, regardless of whether outcomes arose from gambling or choosing the sure option.

Taken together, the EV–CR and RPE magnitude analyses reveal a convergent pattern: SNc stimulation enhances happiness when evaluated relative to the computational structure of the preceding decisions, yet this effect disappears when ratings are averaged across all trials irrespective of context. This dissociation indicates that SNc stimulation does not induce a tonic elevation of momentary happiness, but instead selectively modulates outcome-related happiness. This context-dependent modulation is consistent with DA’s established role in encoding value and prediction error signals, rather than mediating a global increase in baseline happiness.

### SNc stimulation reduces loss aversion and increases risk aversion for losses

We next examined why subjects earned higher rewards following stimulation, using prospect theory (D. Kahneman & Tversky, 2013). The model consists of two stages: computation of subjective utility based on the value of the offered gamble and the safe choice, followed by a decision stage that maps subjective utility to the probability of accepting the gamble. We fit the prospect theory model to choice behavior of our subjects separately during stimulated and non-stimulated trials, followed by comparison of the model parameters (Sokol-Hessner et al., 2009; Tversky et al., 1992). The model separates two distinct aspects of choice. First, risk attitudes describe how people respond to variability in outcomes—for example, whether they prefer a certain outcome or are willing to gamble—and are captured by the curvature of the value function within gains and losses. Second, loss aversion reflects the tendency to weigh losses more strongly than gains of the same size. Importantly, these are dissociable: a person can be risk-averse when choosing between gains, yet still be strongly loss-averse overall. This framework allows us to separately quantify how stimulation affects sensitivity to risk and to losses. Namely, our implementation of the prospect theory model has four key parameters (Figure 4a): the weighting of losses relative to equivalent gains (loss aversion, λ), risk aversion in the gain domain (*α* _*gain*_), risk aversion in the loss domain (*α*_*loss*_), and the degree of randomness in choice behavior (inverse temperature, μ) (see Figure 4a and methods). Models were fit using hierarchical Bayesian inference to the behavior of the subjects (see methods). Model adequacy and parsimony were evaluated with Pareto-smoothed importance-sampling leave-one-out (PSIS-LOO) cross-validation; interpretability was validated with posterior predictive checks and parameter-recovery tests (Aki Vehtari, Gelman, Gabry, & computing, 2017). All model variants fit similarly well and outperform the null models, with good parameter recoverability (see supplementary information).

**Figure 4.**
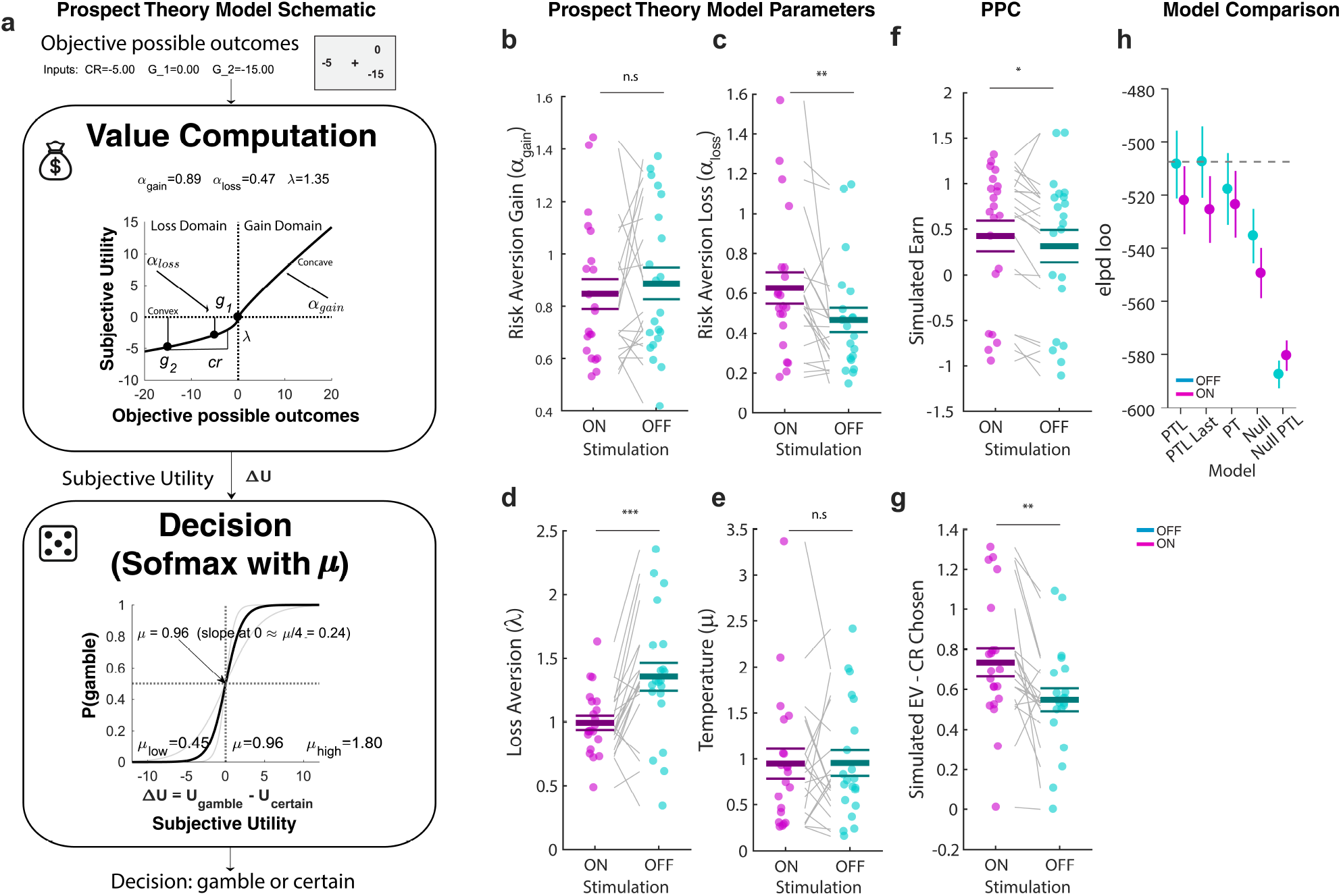
Stimulation effects on Prospect Theory and history-augmented model parameters. (a) Model schematic. Objective outcomes (CR, g1, g2) are transformed to subjective utilities via Prospect Theory with gain and loss curvature (α_gain_, α_loss_) and loss aversion (λ). Choice probability follows a softmax over utility difference, ΔU=Ugamble−Ucertain, with inverse temperature μ (higher μ = less stochastic/steeper choice function). (b–e) Participant-level parameter estimates: α_gain_ (b), α_loss_ (c), λ (d), μ (e). (f–g) Posterior-predictive Check (PPC) simulations: earnings (f) and EV−CR of chosen option (g). (h) Model comparison (ELPD-LOO; higher/less negative indicates better fit). In (b–g), dots are participants; lines connect OFF vs ON; bars show mean ± s.e.m. Paired two-tailed t-tests: *p<0.05, **p<0.01, ***p<0.001; n.s. Colors: stimulation OFF cyan, stimulation ON magenta.

We started by verifying the canonical “diminishing sensitivity” pattern predicted by Prospect Theory—concave value for gains and convex value for losses (Figure 4a, top) —which corresponds to power exponents *α*_*gain*_ and *α*_*loss*_ below one in both domains (Tversky et al., 1992; Wakker, 2010). Consistent with this expectation, across all participants the fitted power exponents were on average less than 1e in both conditions: for gains, α_gain_ < 1 (Figure 4b; ON: *t*_20_ = −2.64, p = 0.01; OFF: *t*_20_ = −1.82, p = 0.083), and for losses, α_loss_ < 1 (Figure 4c; ON: *t*_20_ = −4.72, p = 0.0001; OFF: *t*_20_ = −8.82, p = 2.5 10^−8^). Because α_gain_ < 1 implies concavity of the gain value function, participants were risk-averse in the pure-gain domain (preferring a sure gain over an equal-EV gamble). Conversely, α_loss_ < 1 implies convexity on the loss side. Our subjects thus showed the classic “reflection effect” (D. T. Kahneman, 1979). Importantly, model parameters were not constrained to lie below one, and could in principle take values above one, corresponding to linear or convex value functions.

Comparing the power exponents between ON and OFF stimulation trials revealed that in the presence of stimulation, α_loss_ was larger relative to trials without stimulation (Figure 4c; paired t-test: *t*_20_ = 2.71, p = 0.013), thereby shifting the loss-side exponent toward 1. This upward shift reduces convexity for losses and thereby tempers loss-domain risk-seeking under stimulation (i.e., losses are encoded more linearly, so the subjective “pull” toward taking a loss-domain gamble is reduced). In contrast, α_gain_ showed no significant difference between trials with and without stimulation (Figure 4b; paired t-test: *t*_20_ = −0.76, p = 0.45), indicating that stimulation did not alter risk sensitivity for gains.

We next assessed whether these results could be explained by history effects, i.e. effects of the identity of prior trials. We did so because the stimulation occurred during the outcome period, which could influence the encoding of historical effects. To do so we repeated above analysis for parameter fits derived from two augmented variants of the proposed theory model with additional historical terms (see Methods). In the model with history terms based on previous gamble outcomes, the same qualitative pattern held: both α parameters were less than one in ON and OFF (one-sample tests vs one: α_gain_ ON *t*_20_=-3.50, p=0.002; OFF *t*_20_=-4.65, p=0.0001; α_loss_ ON *t*_20_=-4.08, p=0.0006; OFF *t*_20_=-8.61, p=3.67 10^-08^), and stimulation selectively increased α_loss_ (paired t-test: *t*_20_ = 3.71; p = 0.001) (Supplementary Figure 6a, b). A similar pattern appeared in the second control model, in which the immediately preceding trial’s outcome was included: α_gain_ and α_loss_ were again less than one in both conditions (one-sample tests vs 1: α_gain_ ON *t*_20_= -2.55, p= 0.02; OFF *t*_20_= -3.40, p= 0.003; α_loss_ ON *t*_20_= -4.38, p= 0.0003; OFF *t*_20_= -9.83, p= 4.22 10^-09^), with a selective ON > OFF increase in α_loss_ (paired t-test: *t*_20_ = 4.12; p = 0.0005) and (Supplementary Figure 6i, j). Together, these converging results indicate that stimulation reliably reduced loss-domain risk-seeking by moving the loss curvature toward linearity, while leaving gain-domain risk sensitivity unchanged.

We next turned to examining the loss aversion parameter λ of prospect theory. In general, loss aversion (λ) is expected to be >1 (Kai-Ineman & Tversky, 1979), meaning that losses weigh more than equivalent gains. This was also the case for our subjects: in the stimulation Off condition, λ was significantly higher than 1 (Figure 4d, right; one-sample tests vs 1: *t*_20_ = 3.20; p = 0.004). In contrast, when stimulation was applied the value of λ was not significantly different from 1 (Figure 4d, left; one-sample tests vs 1: *t*_20_ = 0.11; p = 0.91). Thus, patients were more loss averse neutral when being stimulated (Figure 4d; paired t-test ON vs OFF: *t*_20_ = -4.44; p = 0.0002). We again replicated this analysis using the two learning-augmented variants of the model (see supplement).

This model-based characterization is consistent with the model-free analysis shown in Figure 2b,c. There, pooling across stimulation conditions revealed that participants were willing to accept gambles even when they were objectively disadvantageous relative to the certain option (x_50_ < 0), reflecting a baseline tendency toward risk-seeking behavior. Crucially, this effect was specific to the loss domain. When trials were separated by domain (Supplementary Figure 1), a clear negative shift in the indifference point was observed for loss-containing trials (Supplementary Figure 1b,c). Thus, the apparent effect in the pooled analysis is driven by behavior in the loss domain. The prospect theory fits capture this same tendency through convex value functions for losses (α_loss_ < 1), with α_loss_ sufficiently small to overcome the overall loss aversion indicated by λ>1. Importantly, stimulation shifted α_loss_ toward linearity, thereby reducing this convexity and attenuating loss-domain risk-seeking. Together, these results indicate that stimulation did not induce a general change in risk taking but rather normalized a pre-existing bias toward excessive risk seeking in the loss domain.

Our findings provide converging evidence that SN stimulation reduces loss aversion through a combination of lowering λ and increasing α_loss_. Because of the changes in these two parameters, patients’ valuation of losses shifted such that gains and losses were valued more equally.

### SNc stimulation reduces win-stay/loose-shift behavior

After rewards, people tend to repeat their choice (“win–stay”) (Forder & Dyson, 2016; Nowak & Sigmund, 1993; Yajing Zhang, Huynh, & Dyson, 2023), and after non-rewards they tend to avoid making the same choice again (“lose-shift”) (Deng et al., 2016; Donahue, Seo, & Lee, 2013; Forder & Dyson, 2016; Nowak & Sigmund, 1993; Wang, Xu, & Zhou, 2014). In models, this pattern, which in our task would be irrational as the trials are independent, is often captured as an exponential recency weighting of recent outcomes. We next asked whether stimulation modified the effect of choice history on behavior.

In the history-augmented versions of our models, the parameters ϕ_gain_ and ϕ_loss_ capture how recent outcomes modulate subjective utility of the current gambling (see Methods), with positive ϕ_gain_ representing a “win-stay” pattern and negative ϕ_loss_ representing a “lose-shift/avoidance” pattern. Our subjects exhibited both ‘win-stay’ and ‘lose-shift’ behavior, with positive ϕ_gain_ negative ϕ_loss_ in both stimulation ON and OFF trials (one-sample tests vs 0; For ϕ_gain_; ON: *t*_20_ = 3.66; p = 0.002; OFF: *t*_20_ = 9.06; p = 1.62 10^−8^; For ϕ_loss;_ ON: *t*_20_ = −3.85; p = 0.001; OFF: *t*_20_ = −4.12; p = 0.0005; Supplementary Figure 6 d, e, i, m). Both parameters were significantly affected by stimulation, with both ϕ_gain_ (paired t-test: *t*_20_ = −3.68, p = 0.001) and ϕ_loss_ (paired t-test: *t*_20_ = −2.70, p = 0.01) significantly lower in stimulation ON relative to OFF trials. The qualitatively same pattern was also present in the second history-augmented model, which relied only on the immediate prior trial regardless of whether it was a gamble (Supplementary Figure 6 l-m). This data shows that stimulation reduced both win-stay and lose-shift behavior, making decisions less dependent on immediately preceding outcomes. These history effects are present simultaneously with the valuation changes reported for the curvature (α) and loss-weighting (λ) parameters; jointly, the results indicate that SN stimulation in our task during the outcome period both reshaped loss aversion and reduced reliance on win-stay/loose shift strategies (which, in our task, reduce winnings as trials are independent).

### Decision making reliability is lowered by stimulation

We next examined the process that converts subjective utilities into decisions (Figure 4a, bottom). This process is parameterized by the inverse temperature parameter (μ), with higher values indicate more deterministic decisions. In the prospect theory model with no historical terms, stimulation did not significantly change choice stochasticity: μ was not significantly different between ON and OFF (paired t-test: *t*_20_ = −0.05, p = 0.96; Figure 4e). In the models with history terms, a small but reliable difference emerged, with μ lower in ON than OFF trials (paired t-test: *t*_20_ = −2.56, p = 0.02; Supplementary Figure 6 f, n). The absence of an ON/OFF effect in the no-history models suggests μ may partially absorb unmodeled serial dependencies. Once those dependencies are explicitly captured by the φ parameters, a modest decrease in μ under stimulation became detectable, indicating slightly more stochastic choices when stimulation is applied after controlling for history effects.

### Loss-domain curvature explains stimulation-related behavior

The results so far indicate that stimulation affects specific subsets of parameters in the prospect theory model related to how losses are assessed. This raises the question of whether changing a single parameter alone could explain the qualitative change in behavior we found, and if so which one. To examine this question, we next simulated models where we selectively change one parameter at a time.

To follow our reasoning, Figure 5a walks the reader through the four linked steps by which a decision is made in the prospect theory model. The schematic (top-left in Figure 5a, top) shows how offers are evaluated: the certain option (CR) and the two possible gamble outcomes (g1, g2) are mapped from objective outcome values to their corresponding subjective utility via the value function. The gamble’s overall utility is then computed by combining the outcome utilities using U_gamble_ = U(g_1_)/2 + U(g_2_)/2 (see Methods). The difference in utility between the certain option and the gamble are then calculated (ΔU=U_gamble_−U_certain_, middle), followed by conversion to the probability of choosing the gamble (Figure 5a, bottom). CR, g1, and g2 are marked in Figure 5a. The fits shown in Figure 5a are for an example participant, with group average shown in Figure 5b-c.

**Figure 5.**
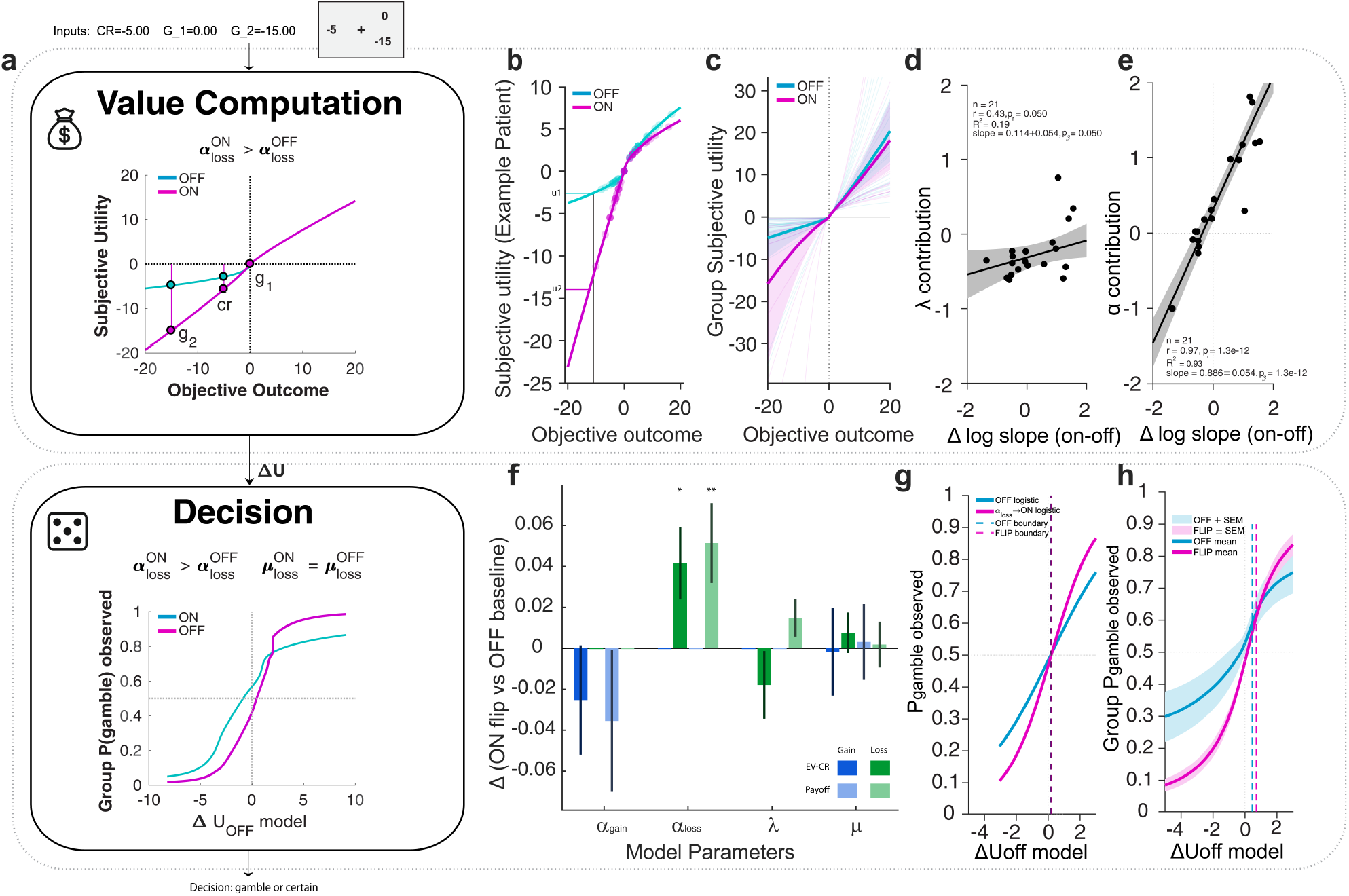
Stimulation-related behavioral changes are best explained by increased loss-domain curvature (α_loss). (a) Prospect Theory + softmax schematic: CR, g1, g2 converted into subjective utilities (separate gain/loss curvature); choices follow a softmax over ΔU=Ugamble−Ucertain. Illustration of the expected effect of higher α_loss_ under ON: the loss-side value function is closer to linear (less convex), increasing the subjective impact of larger losses and shifting ΔU. (b) Example participant subjective-utility curves (OFF vs ON). (c) Group-level subjective-utility functions (mean ± s.e.m.). (d–e) Decomposition of ON– OFF change in the loss-side log-slope at x_0_=−20: the α component accounts for the slope change, whereas the λ contribution is small and not significant (Methods). (f) One-parameter counterfactuals: starting from OFF fits, replace exactly one parameter with its ON estimate and simulate choices on the participant’s trials; only α_loss_ reproduces the ON shift in EV−CR and earnings (mean ± s.e.m.; one-sample two-tailed t-test vs 0). (g–h) Empirical P(gamble) vs OFF-model ΔUOFF (binned; mean ± s.e.m.): ON shows an upward shift that is reproduced by flipping α_loss_ alone (Methods). Colors: stimulation OFF cyan, stimulation ON magenta.

The shape of the curves shows that gains are concave, and losses are convex, consistent with diminishing sensitivity, with stimulation making the loss side less convex and less compressed over the task-relevant loss range, as expected from a larger α_loss_ in the stimulation condition. Importantly, in our model the relative weighting of equal-magnitude losses and gains is not determined by λ alone, because gains and losses have separate curvature parameters. Specifically, |U(−x)|/U(x)=λx^αloss−αgain^, so the apparent loss sensitivity of the full utility curve depends jointly on λ, α_loss_, α_gain_, and outcome magnitude. Therefore, a reduction in the fitted loss-aversion parameter λ can coexist with a less compressed and locally steeper loss branch when α_loss_ increases.

To examine how much of the stimulation-triggered change in the loss-side value-function slope comes from loss weight λ vs. loss curvature α_loss_, we analytically decomposed the change in log slope at a fixed loss of L =-20 magnitude (see Methods). Empirically, the α_loss_ contribution alone tracked the total slope change observed in our subjects very well (Figure 5e; slope =0.886±0.054, R^2^=0.932, r=0.97, p=1.3 10^−12^, n=21), whereas the λ contribution showed only a weaker association (Figure 5d; slope =0.114±0.054, R^2^=0.192, r=0.43, p=0.050, n=21). Repeating the same analysis at different loss magnitudes leads to similar results (i.e., α_loss_ L = -5, slope =0.816±0.071, R^2^=0.88, r=0.94, p=5 10^−10^, n=21; λ slope = 0.184±0.071, R^2^=0.26, r=0.51, p=0.018, n=21). This analysis supports the hypothesis that it is the curvature as indexed by the value of α_loss_, rather than the overall loss weight λ, as the primary parameter that drives the stimulation effects we found.

We next tested this hypothesis using a simulation. For each subject, we used the model to generate choices for the same stimuli that were shown to the subject. The parameters of the model were first all set to those estimated from the stimulation OFF condition (‘baseline’), followed by selectively changing only a single parameter at a time to the value estimated from the stimulation ON condition. We then compared the (i) expected payoff and (ii) probability-weighted EV–CR for each trial between the baseline simulation and the simulation where only a single parameter was changed (see Methods). This approach revealed that the only single-parameter change that reproduced the empirical data was that of α_loss_, for which both the changes in EV-CR and earnings/trial were positive as observed in the data (Figure 5f; Δ(EV–CR |accepted, loss trials one-sample tests vs 0: *t*_20_ = 2.35; p = 0.03; Δ(Earnings/trial, loss trials one-sample tests vs 0: *t*_20_ = 2.63; p = 0.016). In contrast, modifying α_gain_, λ, or μ yielded differences that were not significantly different from zero (Figure 5f, all p > 0.05). This suggests that increasing loss-side curvature by modifying α_loss_ is sufficient to account for the change in behavior we observed that results from SN stimulation (Figure 5f). To illustrate this effect, we re-plotted the subjective utility curves with all parameters set the stimulation OFF value except that of α_loss_, which was set to the value estimated from the stimulation ON trials. In both an example subject (Figure 5g) and the group mean (Figure 5h), this single parameter change modified the utility function in the qualitatively same manner than the stimulation did. The decision boundary (Figure 5g-h, dashed lines), in contrast, remained essentially identical.

Together, these results indicate that stimulation improves risky decision making primarily by altering loss-domain curvature: increasing α_loss_ reduces compression of larger losses and increases the local slope of the loss branch over the task-relevant range. Consequently, subjects reject more loss-heavy gambles, which in turn increases overall payout. These findings indicate that dopaminergic stimulation reduced loss-domain risk seeking, improving expected-value efficiency and earnings.

## Discussion

In our task, participants exhibited a baseline tendency toward risk-seeking in the loss domain, accepting gambles even when they were objectively disadvantageous relative to the certain option. Focal microstimulation of the human SNc reduced this bias, shifting behavior toward greater alignment with objective value and resulting in improved earnings. This improvement was driven by more selective acceptance of advantageous gambles (higher EV–CR among chosen options), rather than systematic changes in outcomes when participants chose the certain option. These findings indicate that stimulation did not globally increase risk taking but instead normalized a pre-existing bias toward disadvantageous “loss-chasing” choices. Such value-selective effects are consistent with distributed valuation-network accounts implicating ventral striatum and vmPFC interactions with midbrain dopaminergic systems in value-based decision making (Bartra et al., 2013; Haber & Knutson, 2010; Kable & Glimcher, 2009).

A key motivation for our study was to move beyond correlational evidence and ask what aspects of decision computations can be influenced by causally manipulating midbrain activity. By stimulating the SNc in this manner, we show that the SNc plays a causal role in the computations that determine choice in a task that does not involve learning. This is in contrast to most prior mechanistic work on the dopaminergic midbrain, which has emphasized its role in learning rather than in decision making (D’Ardenne et al., 2008; Kishida et al., 2016; Schultz et al., 1997; Steinberg et al., 2013; Tsai et al., 2009). It has long been known that dopaminergic signals encode variables relevant to economic decision making such as utility and marginal utility (Schultz, 2016a; Stauffer, Lak, & Schultz, 2014), which suggests a role of these signals in decision making rather than (or in addition to) learning. Here, we now show causal evidence using microstimulation.

We note that our causal manipulation involved microstimulation through a high-impedance microelectrode (30 μA current). Such stimulation is highly focal and is thought to modulate only a small population of neurons in the immediate vicinity of the electrode tip. Classic microstimulation studies in non-human primates have demonstrated that currents in this range can influence behavior by perturbing the activity of relatively small neuronal ensembles. For example, stimulation in area MT was estimated to affect only tens to at most a few hundred neurons yet was sufficient to reliably bias perceptual decisions about motion direction (Salzman, Britten, & Newsome, 1990; Salzman, Murasugi, Britten, & Newsome, 1992; Tehovnik, Tolias, Sultan, Slocum, & Logothetis, 2006). Our approach therefore aligns with this tradition of causal circuit perturbation, where subtle manipulations of localized neuronal populations can produce measurable changes in behavior.

This approach differs fundamentally from clinical (DBS), which delivers currents in the milliampere range through large, low-impedance electrodes and thus affects a substantially larger volume of tissue. DBS is therefore thought to modulate distributed networks rather than small neuronal ensembles near the electrode tip. The present results suggest that modulating a very small population of neurons within the human SNc is sufficient to alter decision making under risk and the affective evaluation of outcomes. This sensitivity highlights the potential importance of localized dopaminergic signaling in shaping value-based behavior and momentary affect.

These findings may also help contextualize behavioral changes reported in patients receiving clinical DBS. In STN DBS has been associated with alterations in reward processing and impulse control, including pathological gambling and increased risk-taking in some PD patients (Frank, Samanta, Moustafa, & Sherman, 2007; Jahanshahi, Obeso, Baunez, Alegre, & Krack, 2015; Voon et al., 2006). Our results suggest that even highly focal modulation of dopaminergic neurons in the human SNc can influence value-based decisions and affective responses to outcomes, raising the possibility that DBS-induced behavioral changes may partly arise from perturbations of dopaminergic circuits involved in valuation and reward prediction.

Critically, the effect of stimulation on behavior in our task was not a global increase in risk taking, but rather a reduction of a pre-existing tendency toward risk-seeking in the loss domain. This distinction matters because systemic dopaminergic manipulations in humans, such as that which results after the administration of levodopa in combination with a peripheral decarboxylase inhibitor (e.g., benserazide), can increase gambling propensity without improving payoff. For example, in a closely related mixed-gamble/happiness paradigm, levodopa administration increased risk-taking and altered affective responses without reliably increasing overall earnings (Rutledge et al., 2015). Related work suggests that dopamine can increase value-independent gambling propensity under some pharmacological manipulations (Rigoli et al., 2016). In contrast, our stimulation effects were value-selective: choices shifted toward offers with more favorable EV –CR, consistent with increased expected-value efficiency rather than a uniform elevation of gambling.

Our computational modeling grounds these behavioral effects in specific prospect theory components. We used a canonical from of Prospect Theory (D. T. Kahneman, 1979; Tversky et al., 1992; Wakker, 2010) that has been used extensively to model behavior during the kinds of mixed gambles our subjects performed (Rutledge et al., 2015; Sokol-Hessner et al., 2009) to examine how stimulation affected decision making computations. We find that SNc stimulation jointly reduced loss aversion (λ shifted toward 1) and attenuated the canonical risk-seeking curvature for losses (α_loss_ increased toward linearity), while leaving gain curvature and choice stochasticity comparatively unchanged. Although reduced λ and reduced loss-domain risk seeking can seem superficially contradictory, these parameters capture dissociable computations: λ scales the relative weight of losses versus gains, whereas α_loss_ shapes sensitivity within the loss domain and thus the temptation to accept disadvantageous “loss-chasing” gambles. Under stimulation, losses were globally less overweighted (lower λ), while large prospective losses were no longer psychologically underweighted by convexity (α_loss_ closer to 1), producing a combination that favors rejecting strongly disadvantageous offers but preserves acceptance of objectively favorable mixed gambles.

This mechanistic interpretation is consistent with neurophysiological evidence that dopaminergic signals can encode subjective value and marginal utility rather than raw outcome magnitude (Schultz, 2016a; Stauffer et al., 2014), which provides a plausible link between SNc perturbation and selective changes in loss-side curvature. Importantly, the valuation-stage specificity implied by the parameter pattern also aligns with distributed-valuation accounts in which vmPFC/ventral striatum instantiate subjective value representations that can be influenced by dopaminergic inputs (Bartra et al., 2013; Haber & Knutson, 2010; Kable & Glimcher, 2009).

Beyond static valuation, the history-augmented results suggest that SNc stimulation reduced short-lived outcome-history biases (win-stay/lose-shift) that are maladaptive in our task because outcomes are independent across trials. We hypothesize this is the case because our stimulation disrupted outcome signals, thereby reducing history dependence. Dopaminergic state is well known to modulate how outcomes shape subsequent behavior (Frank, Seeberger, & O’reilly, 2004; Pessiglione, Seymour, Flandin, Dolan, & Frith, 2006), and dopaminergic manipulations in related paradigms can alter both choice and affective dynamics (Rutledge et al., 2015). In our data, stimulation pushed choices toward greater reliance on value differences (EV–CR / ΔU) and away from noisy carry-over from prior wins/losses—consistent with a shift toward more normatively efficient “myopic” value-based choice in an environment where myopia is optimal.

A second motivation for conducting the present study was to examine the relationship between SNc activity and subjective well-being. We found that SNc stimulation did not produce a uniform elevation of subjective well-being (i.e. happiness). Instead, it modulated happiness in a context-dependent manner, enhancing outcome-related happiness when evaluated relative to the computational properties of preceding decisions. Although the strongest condition-wise effects were observed following positive outcomes in advantageous contexts (e.g., high EV–CR and high-RPE wins), these were not accompanied by reliable interactions, indicating that stimulation broadly amplifies outcome-related happiness rather than acting exclusively on specific trial types. This pattern suggests a strong influence of expectations on the hedonic impact of outcomes, as shown in our analysis by the significant main effects of EV-CR and RPE. The findings dovetail with computational accounts in which momentary happiness reflects recent expectations and prediction errors rather than cumulative earnings (Rutledge et al., 2014), and with the broader view that dopaminergic signals incorporate internal decision variables such as belief in choice accuracy or confidence (Lak, Nomoto, Keramati, Sakagami, & Kepecs, 2017). More broadly, this context-dependent coupling of outcomes to experienced happiness supports the idea that “more reward” does not map linearly onto “more happiness” in humans (D. Kahneman & Deaton, 2010), highlighting the importance of mechanistic interventions that can alter how decision context shapes affect. Notably, our findings differ from those in a study utilizing L-DOPA (Rutledge et al., 2015), which found that tonic dopaminergic enhancement increased gambling probability and selectively boosted happiness after small rewards from low-value gain gambles without improving overall earnings. In our data, in contrast, SNc stimulation neither indiscriminately increases risk-taking nor uniformly elevates happiness; instead, it enhances outcome-related affect in a manner that aligns subjective experience with decision quality.

Finally, our findings are of relevance to clinically observed dopamine-sensitive alterations in valuation and risky behavior in patients with PD. Medicated PD patients can show reduced loss aversion in effort-based decision contexts (Chen et al., 2020), and impulse-control disorders including pathological gambling are elevated in PD populations treated with dopaminergic medications (Heiden et al., 2017; Santangelo et al., 2013). Moreover, neuromodulation within basal ganglia circuits has been associated with changes in risky behavior, including reports of pathological gambling after STN DBS (Smeding et al., 2007) as well as broader evidence that DBS can modulate risk-taking depending on task structure (Brandt et al., 2015). Our parameter-level results suggest one candidate computational route through which focal basal ganglia/midbrain perturbations could shift risky choice: concurrent changes in loss weighting (λ) and loss curvature (α_loss_), together with reduced reliance on recent outcome history, may move behavior toward greater expected-value efficiency in some contexts while still allowing clinically relevant shifts in risky behavior in others. Our results highlight the need to better understand (and avoid) the causal effect of STN DBS on activity in the SNc, which given the proximity of the STN and the connectivity from the STN to the SN is expected to be prominent in many instances. Our findings suggest that our task, and fitting of the prospect theory model to the behavior during, offers a sensitive new approach to rigorously investigate the effects of different kinds of STN stimulation on risky decision making and to assess potential strategies to reduce the cognitive deficits that can result from PD and/or the treatment of PD symptoms with STN.

Our conclusions are supported by model robustness checks and model comparison. Across prospect theory variants, predictive performance was similar and systematically better than null alternatives, consistent with the view that the key ON–OFF differences reflect reliable features of the choice process rather than overfitting. Our use of PSIS-LOO for Bayesian model evaluation follows established best practices for comparing Bayesian models using posterior draws (Aki Vehtari et al., 2017; A Vehtari, Simpson, & Gelman, 2017).

A key consideration in interpreting our findings is the extent to which ongoing neurodegeneration in PD may shape the properties of the dopaminergic signals we measured. PD is characterized by a progressive and regionally heterogeneous loss of dopamine neurons within the substantia nigra, with the most severe degeneration typically observed in ventral and caudal tiers, while more dorsal regions are relatively spared, particularly at earlier disease stages (Damier et al., 1999; Fearnley & Lees, 1991). Given that our recordings were obtained during STN-targeted DBS procedures, sampled neurons were predominantly located in these more dorsal regions, where a substantial population of dopaminergic neurons remains functional. Consistent with this, prior human electrophysiological studies have successfully recorded putative dopaminergic neurons in the substantia nigra of PD patients undergoing DBS surgery, demonstrating preserved encoding of reward, value, and contextual signals by putative DA neurons (Batten et al., 2024; Imtiaz et al., 2026; Imtiaz et al., 2024; Ramayya et al., 2014; Zaghloul et al., 2009). The patients included in this study were at relatively early stages of disease compared to post-mortem cohorts, suggesting that a substantial proportion of dopaminergic neurons remained viable in the regions sampled. Our recordings confirmed this interpretation, and we excluded the subjects in whom we did not identify at least one putative DA neuron as a pre-caution. Together, these considerations support the interpretation that the effects we observe reflect the activity of a functionally meaningful dopaminergic population, while also highlighting that the precise relationship between disease progression and dopaminergic signaling in humans remains an important question for future work.

## Conclusions

In summary, focal microstimulation of the human SNc was sufficient to reshape both decision making under risk and subjective happiness. In our task, participants exhibited a baseline tendency toward risk-seeking in the loss domain, accepting gambles even when they were objectively disadvantageous relative to the certain option. SNc stimulation reduced this bias, shifting behavior toward greater alignment with objective value. Rather than globally increasing gambling or happiness, stimulation improved the quality of risky choices, increased earnings, and reduced maladaptive dependence on recent outcomes. In parallel, stimulation did not tonically elevate momentary happiness, but instead modulated affect in a context-dependent manner, enhancing happiness in relation to decision outcomes and their associated value signals. Computationally, these effects were best explained by altered loss processing, particularly a reshaping of the loss function that reduced loss-domain distortion. Together, these findings provide causal evidence that localized perturbation of human midbrain dopaminergic circuits can influence valuation, choice, and momentary well-being, offering a mechanistic bridge between dopamine’s role in value and prediction error signaling and its impact on adaptive behavior and subjective experience.

## Methods

### Patients and Electrophysiological recordings

All subjects in this study were tested while undergoing surgery for the implantation of a DBS device for treatment of PD. A total of 24 subjects volunteered for this study (see Supplementary Table 1 for demographics; age at the time of recording was 68.5 ± 7.85 years, UPDRS III score off medication, pre-DBS treatment: 21.9 ± 11.12; n = 7 females). Subjects performed 24 sessions in total. Subjects were off their PD medication for at least 12 hours prior to surgery. All subjects provided informed consent (Feinsinger et al., 2022), and all protocols were approved by the Institutional Review Board of Cedars-Sinai Medical Center.

Identifying the clinical target, the STN, involves preoperative imaging and intraoperative microelectrode recordings. Indicators of STN entry include increased background neuronal activity, multiunit large spike activity, and responses to limb movements. We marked the ventral STN border by a drop in background activity, followed by higher frequency and more regular activity in the substantia nigra reticulata (SNr). However, in the ventral STN, slower firing rates and atypical patterns, along with lower neuron density, can make precise identification challenging, as SNr neurons may exhibit similar characteristics to some STN neurons. To aid in targeting the ventral border of the subthalamic nucleus, we also utilized microstimulation induced inhibition as suggested in (Lafreniere-Roula, Hutchison, Lozano, Hodaie, & Dostrovsky, 2009; Lafreniere-Roula et al., 2010). Their findings indicate that prolonged inhibition of firing following low-amplitude, high-frequency microstimulation via the recording electrode is a consistent feature of nearly all substantia nigra (SN) neurons and rarely, if ever, occurs in STN neurons (Figure 1e).

Single neuron activity was recorded during anatomical mapping while surgically implanting a Deep Brain Stimulation (DBS) device (for details on surgery and target mapping, see (Kamiński et al., 2018b)). Micro-electrodes (Alpha Omega microelectrodes, STR-009080-00) used had a micro-wire at the tip and a larger macro-contact 3 mm above it. In each session, one microelectrode was placed along the target trajectory, while a second electrode was sometimes positioned in a Bengun array either 2 mm medial/lateral or anterior/posterior. The electrodes were grounded and referenced to guide cannulae located 25 mm above the target site. Broadband signals (2Hz-44 kHz) were recorded using a NeuroOmega system (Alpha Omega Inc.). Both microelectrodes were advanced simultaneously in 1 mm steps, monitoring single neuron activity and comparing it to the expected trajectory based on surgical imaging. As the electrodes approached the STN, the advancement speed was reduced to 0.1-0.5 mm per step to go 3mm below target towards SN, and single neurons were carefully monitored. Once single unit spiking was detected on at least one microelectrode, we waited at least one minute to ensure stable spike amplitude before starting the experiment.

### Electrode localization

Precise localization of intracranial microelectrodes was performed following the protocol in Mosher et *al*., 2021. Post-operative computed tomography (CT) scans, which clearly visualized the clinical DBS electrodes, were aligned (BRAINSFit 6DOF; (Johnson, Harris, & Williams, 2007)) with high-resolution pre-operative magnetic resonance imaging (MRI) scans with detailed anatomical context. Alignment quality was visually verified.

Next, the tip of the DBS electrode was marked manually on the CT scan and traced along each slice of the axial CT (3D Slicer 5.0.3). This trace defined the trajectory of the DBS electrode and was used to infer the location of the microelectrode recording in native patient space, as in Mosher et *al*., 2021.

To obtain standardized coordinates for the microelectrode locations, we next calculated a sequence of transformations that warped the structural MRI to a template brain in Montreal Neurological Institute (MNI) space (Fonov, Evans, McKinstry, Almli, & Collins, 2009). This transformation included both affine and nonlinear warping components (FreeSurfer 7.4.1, Advanced Normalization Tools ANTs) to account for global differences in brain orientation and local anatomical variability across individuals. The transformations were then applied to the microelectrode locations—originally defined in native space—to map them to the standardized MNI space.

The resulting microelectrode electrode locations in MNI space were used for group-level analyses, anatomical labeling, and spatial visualization. To visualize electrode locations across subjects, we employed two complementary methods. First, electrodes were projected into 2D using the Distal Atlas (Ewert et al., 2018) to provide an overview of anatomical distribution in standardized coronal and axial planes. Second, we reconstructed electrode positions using the Lead-DBS toolbox, visualized in relation to the STN atlas developed by (Zhang, Larcher, Misic, & Dagher, 2017). These 3D renderings, shown from two viewing angles, facilitated an anatomically detailed and spatially intuitive representation of recording locations across the patient cohort.

### Task

We used a similar version of a probabilistic gambling task used in (Rutledge et al., 2014). Stimuli were presented using MATLAB (MathWorks, Inc.) and Psychtoolbox. Patients performed up to 120 trials per session. Each trial required subjects to choose between a certain option and a gamble, with both options having equal probabilities for two outcomes (Figure 1a). There were three trial types: mixed trials (a certain amount of 0 vs. a gamble with equal probabilities of a gain or a loss), gain trials (a certain gain vs. a gamble with a 0 or a larger gain), and loss trials (a certain loss vs. a gamble with a 0 or a larger loss) (Figure 1b). Choices remained on the screen for 0.9 seconds before the outcome was displayed for 1 second, followed by a 1.1-second intertrial interval. Each trial type was repeated 20 times, resulting in a total of 120 trials.

We used three different trial types (Figure 1b). Of the 120 choice trials, there were 40 mixed trials, 40 gain trails, and 40 loss trials. The mixed trials involved a choice between a certain 0 and a gamble with equal chances of a monetary gain or loss. There were four possible gain amounts (300, 500, 800, 1100) and loss amounts determined by 5 multipliers (0.2, 0.4, 0.52, 0.82, 1.2), to vary loss sensitivity. The 40 gain trials involved a choice between a certain gain and a gamble with equal chances of a larger gain or 0. There were four certain amounts (200, 300, 400, 500) and the gamble gain was determined using 5 multipliers on the certain amount (1.68, 2, 2.48, 3.16, 4.2). The 40 loss trials used the same monetary amounts as the gain trials, with four certain amounts (−200, −300, −400, −500) and the same 10 multipliers as for the gain trials. The maximum gamble gain or loss per trial was 2100.

After every three trials, subjects were asked, “How happy are you at this moment?” Following a 1.1-second delay, a rating line appeared, and subjects had 4 seconds to move a cursor along the scale using button presses. The scale ranged from “very unhappy” on the left end to “very happy” on the right end, with the cursor starting at the midpoint. Each rating was followed by a 1.1-second intertrial interval. Subjects completed 120 choice trials and answered the happiness question 40 times. All responses were given by moving the ball of a big track ball mouse (ErgoGuys 12000006 Ablenet Bigtrack Trackball Mouse) to the right ot to the left.

### Trial balancing between stimulation conditions

We used random subsampling of trials to assure that trials were appropriately balanced between the stimulus ON and OFF conditions despite variability in behavior. To do so, we constructed a balanced subset of trials that are matched 1:1 across conditions on the objective inputs (CR, g1, g2) and the realized outcome index s, up to a tolerance of τ=10 in task units.

Let each trial t be encoded by:

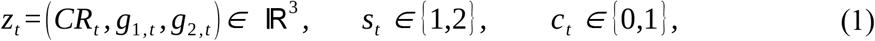

where *CR*_*t*_ is the certain reward, *g*_1, *t*_ and *g*_2,*t*_ are the two gamble outcomes, *s*_*t*_ indicates which outcome was realized on that trial (1→*g*_1_, 2→*g*_2_), and *c*_*t*_ is the condition (0=OFF, 1=ON).

To compute the distance and caliper, for a candidate ON–OFF pair (i, j) with ci=1 and cj=0, we defined a Euclidean distance over the continuous inputs:

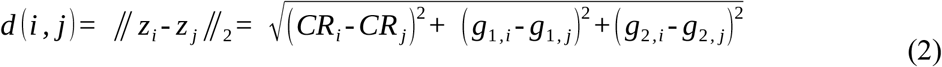

Pairs were admissible only if they satisfied an exact match on the realized outcome and a caliper on distance:

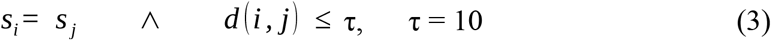

We used a matching algorithm (1:1 without replacement). We performed nearest-neighbor matching without replacement from ON to OFF. We first randomly permute the ON trials to avoid order bias (fixed seed for reproducibility). We then for each ON trial *i* in that order, find the OFF trial *j* with smallest *d* (*i*, *j*) among those not yet used that also satisfy *s*_*i*_ = *s*_*j*_ and *d* (*i*, *j*) ≤ τ. And lastly if such *j* exists, form the pair (*i*, *j*) and remove *j* from further consideration; otherwise *i* is left unmatched (and excluded). This yields a set of pairs 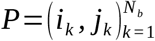. The balanced dataset consisted of the union of these matched ON and OFF trials; all unmatched trials were excluded. In Supplementary Figure 2, matched trials are labeled kept and unmatched trials removed. We compute the balance diagnostics to verify balance between conditions after matching by (i) QQ-plots and empirical CDF overlays for EV, Outcome, RPE, EV−CR, and CR; and (ii) two-sample *t* and Kolmogorov–Smirnov tests on the trial-level distributions (and, where reported, paired t tests on participant means). These analyses confirmed that stimulus features were statistically indistinguishable between ON and OFF in the balanced set, ensuring that downstream ON–OFF behavioral differences cannot be attributed to differences in the presented stimuli (Supplementary Figure 2).

### Stimulation artefact removal

We used a semi-automated custom zero padding algorithm in order to remove the stimulation artefact from the recorded signal. We first down sampled the recorded signal from 44 to 32kHz. We then detected peaks higher than 100 standard deviations from the mean to select the highest stimulation pulses artifacts normally seen at the end of stimulation trains and zero padded from 50 before to 300 samples after the detected peaks (Supplementary Figure 2a). Next, we repeated this step detecting peaks higher than 50 standard deviations from the mean to remove remaining stimulation artefacts that were found in the signal normally at the beginning of the stimulation trains. Then, we visually inspected each stimulation train and set a custom threshold to find the stimulation artefacts. We used between 3 and 6 standard deviations from the mean and zero padded 10 samples before and after the detected peaks (Supplementary Figure 2b). We then inspected visually the beginning of each stimulation train, since we found the bigger peaks exhibited a longer rebound in the recorded signal. When those stimulation artefacts exceeded 40 standard deviations, we removed from 50 samples before the peak to 200 samples after. On average the total percentage of removed signal is 0.81% ± 0.26% (mean ± std) (Supplementary Figure 2c).

### Spike sorting and quality measures

To detect and sort spikes from putative single neurons in each wire we used the semiautomated template-matching algorithm Osort (Fried, Rutishauser, Cerf, & Kreiman, 2014; Rutishauser, Schuman, & Mamelak, 2006). Spikes were detected after bandpass filtering the raw down sampled masked signal in the 300-3,000 Hz band. Quality measures were computed for all the sorted putative single neuron to measure the recordings and sorting quality (Harris, Henze, Csicsvari, Hirase, & Buzsaki, 2000; Kamiński et al., 2017; Pouzat, Mazor, & Laurent, 2002) (see Supplementary Figure 3 for single cell quality metrics). We computed the percentage of interspike intervals (ISIs) below 3 ms, the ratio between the s.d. of the noise and the peak amplitude of the mean waveform of each cluster (peak SNR), the pairwise projection distance in clustering space between all neurons isolated on the same wire (projection test; in units of s.d. of the signal) (Pouzat et al., 2002); the modified coefficient of variation of variability in the ISI (CV2). All analysis in this paper is based on signals recorded from micro wires. We ensured that there was no stimulation artefact specifically looking for unexpected frequency peaks in the power spectrum at the frequency of the stimulation train 130Hz and harmonics.

### Identifying putative DA and Gaba activity neurons

Electrophysiology studies in non-human primates typically identify presumed DA and GABA units based on the characteristics of extracellular spike waveforms recorded by microelectrodes (Fiorillo, Yun, & Song, 2013). Previous research combining electrophysiological recordings with pharmacological manipulations (Schultz & Romo, 1987) or histochemical techniques (Henny et al., 2012) has shown that DA neurons exhibit low firing rates and broad waveforms, whereas GABA neurons show fast firing rates and narrow waveforms (Ungless & Grace, 2012). For each unit, we estimated the baseline firing rate by calculating the mean firing rate over the entire recording session and measured the waveform duration by assessing the peak-to-trough duration (Barto, Singh, & Chentanez, 2004). We identified putative DA units as those with baseline firing rates slower than 15Hz and waveform durations greater than 0.5 ms, and GABA units as those with baseline firing rates faster than 15Hz and waveform durations less than 0.5 ms. Similar parameters were used in (Barthó et al., 2004; Ramayya et al., 2014).

### Behavioral analyses

In order to analyze the behavioral data, we separated trials into stimulation ON and OFF. Since the stimulation was deliver during the outcome period of a subset of trials, we consider in the behavioral analysis as stimulation trials those that followed a stimulated trial. For each trial EV was computed as the average between the two offered gambles, CR as the quantity offered and RPE as the reward minus the EV. If the CR was chosen on a given trial, EV and RPE of that trial was set to 0. If the gamble was chosen in a trial, CR was set to 0 for that trial. Thus, in Figure 2d total earnings are shown collapsing trials where the gamble and certain reward was chosen, only when gamble was chosen in Figure 2f, and when certain option was chosen in Figure 2g. EV-CR in Figure 2h and i are shown for trials where gamble was chosen. The distance between certain reward and the expected value for the gamble was computed as a metric for the gamble worth or objective outcome difference. If CR-EV>0 (CR>EV) the certain would be better option whereas if CR-EV>0 (CR<EV) the gamble would be more worth it. Throughout the manuscript, we use one sample, paired t-tests, ANOVAs, or mixed-effects GLMs (fitglme in MATLAB) to assess statistical differences between conditions.

### Psychometric analysis and estimation of indifference point (x_50_)

To characterize choice behavior, we constructed trial-wise predictors based on the difference between the expected value of the gamble and the certain option (EV−CR). Mixed (gain–loss) and pure-loss trials were combined to form a pooled “loss-containing” set of trials. Choices were coded as binary (gamble vs certain). A logistic regression model with subject-specific intercepts was fitted to the pooled data (ON and OFF) to estimate the probability of choosing the gamble as a function of EV−CR.

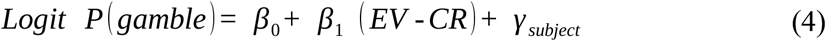

Population-level psychometric curves were obtained by averaging model predictions across subjects. The indifference point (x_50_) was defined as the value of EV−CR at which the predicted probability of gambling was 0.5. Negative x_50_ values indicate a bias toward gambling even when the expected value of the gamble is lower than the certain option, consistent with loss-seeking behavior.

Uncertainty in x_50_ was quantified using a bootstrap procedure in which subjects were resampled with replacement and trials were resampled within subject. For each bootstrap sample, the model was refit and x_50_ recomputed. Statistical significance was assessed using a one-sided bootstrap test evaluating the proportion of bootstrap estimates greater than or equal to zero (testing x_50_ < 0). Confidence intervals were computed as the 2.5th and 97.5th percentiles of the bootstrap distribution.

### Happiness analysis

On a subset of trials, participants provided a happiness rating reflecting their current momentary well-being. To test whether stimulation modulated the hedonic impact of outcomes as a function of decision quality, we adapted the model-free approach introduced by (Rutledge et al., 2015; Rutledge et al., 2014).

For each happiness rating, we first identified the most recent preceding trial on which the participant had made a choice (sure option or gamble). For that preceding choice, we computed the decision value EV-CR = EV_gamble_−CR, where EV_gamble_ is the mean of the two possible gamble outcomes and CR is the certain amount available on that trial. This EV–CR term indexes how advantageous the offered gamble was relative to the sure option, independent of which option was chosen.

Within each participant and stimulation condition (ON/OFF), EV–CR values associated with ratings were split by the within-subject median into low-value and high-value decision contexts. Outcome valence for the preceding choice was coded as win (non-negative realized outcome) or loss (negative outcome). Each happiness rating was thus assigned to one of four cells, separately for ON and OFF: Low EV–CR win, Low EV–CR loss, High EV–CR win, High EV–CR loss.

For the primary (all-trials) analysis, the “last chosen option” could be either the sure option or the gamble; EV–CR was always computed from the corresponding trial’s offer. In a secondary, gamble-restricted analysis, we repeated the same procedure considering only ratings linked to trials on which the gamble had been chosen, to align more directly with prior work.

For each participant and condition, we computed the mean happiness in each EV–CR × outcome and compared ON vs. OFF using paired-sample tests (see Results). This framework allowed us to assess whether SN stimulation produces a global shift in mood or a selective change in the hedonic impact of outcomes in more vs. less advantageous decision contexts, without relying on parameters from the Prospect Theory models.

### Happiness Computational Model

We modeled intermittent happiness ratings using a Rutledge-style linear model (Rutledge et al., 2014) in which ratings were explained by exponentially decayed histories of task variables. Trials were indexed by *t* = 1, *…, T*. On each trial, the certain reward was *CR*_*t*_ (the sure option), the gamble expected value was *EV*_*t*_*=(G1*_*t*_*+G2*_*t*_*)/2* (mean of the two gamble outcomes), and the realized outcome of the chosen option was *OUT*_*t*_ defined as *OUT*_*t*_ *= CR*_*t*_ for sure choices and *OUT*_*t*_ equal to the realized gamble outcome for gamble choices. Reward prediction error was defined as:

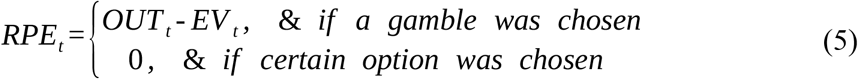

To capture integration of recent experience, we constructed exponentially decayed state variables for each regressor *xt*∈*{CR*_*t*_,*EV*_*t*_,*RPE*_*t*_*}* using a single decay parameter γ∈[0,1):

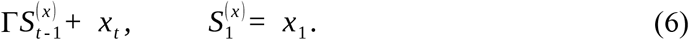

Let Ɍ denote the set of trials at which a happiness rating was provided and let *H*_*t*_ be the participant’s rating at *t*∈ Ɍ (z-scored within participant). We modeled happiness ratings as

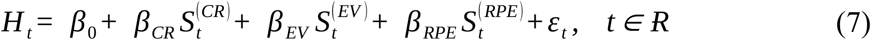

For a fixed γ, coefficients were estimated per participant using ridge regression with penalty λ=10^−2^ applied to non-intercept coefficients (the intercept was not penalized). The decay parameter was selected by grid search. For the primary in-sample analyses we searched *γ*∈ *{0,0*.*05,…,0*.*95}*; for the condition-specific (stimulation OFF/ON) fits we used a finer grid *γ* ∈ *{0,0*.*01,…,0*.*99}*. Model fit at rating times was summarized by the Pearson correlation *r* between fitted and observed ratings and by the coefficient of determination *R*^2^. Group-level significance of model fit was assessed with a one-sample *t*-test on Fisher z-transformed correlations z=tanh^−1^(r). To test whether fits exceeded chance, we generated a within-participant null distribution by randomly permuting the trial-wise *RPE*_*t*_ sequence, recomputing 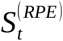, and refitting the model (including re-optimizing γ) for each shuffle (200 shuffles; random seed 1).

### Parametric decision model based on Prospect Theory

We modeled choice behavior using prospect theory (D. Kahneman & Tversky, 2013), as described by (Sokol-Hessner et al., 2009) and later applied by (Rutledge et al., 2015).

To fit a prospect theory model, we used Stan, a probabilistic programming language for Bayesian inference, and interfaced it through Python using the CmdStanPy/PyStan library. The model was specified using a hierarchical Bayesian approach, incorporating subject-level parameters.

We implemented the model using Stan’s Hamiltonian Monte Carlo (HMC) sampling, running it through Python. The model was written in Stan’s modeling language, defining the likelihood function for choices based on prospect theory.

This model characterizes choice behavior using four key parameters. The subjective values, or expected utilities, of gambles and certain options were calculated with the following equations:

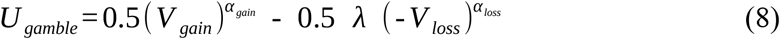

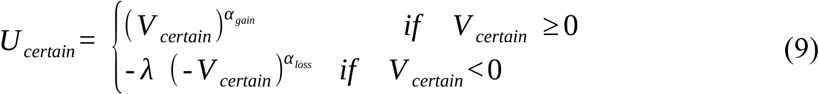

Here, *V*_*gain*_ and *V*_*loss*_ represent the values of the potential gain and loss from a gamble, respectively, while *V*_*certain*_ represents the value of the certain option. In gain trials, potential losses were set to zero (*V*_*loss*_ = 0), and in loss trials, potential gains were set to zero (*V*_*gain*_ = 0). The curvature of the utility function in the gain domain, determined by *α*_*gain*_, reflects the degree of risk aversion. A risk-neutral individual in gain trials has *α*_*gain*_ = 1, indicating indifference between a certain gain and a gamble with the same expected value. A risk-seeking individual would have *α*_*gain*_ > 1, while a risk-averse individual would have *α*_*gain*_ < 1. Similarly, *α*_*loss*_ determines risk aversion in the loss domain, with risk-seeking individuals having *α*_*loss*_ < 1 and risk-averse individuals having *α*_*loss*_ > 1.The parameter λ measures loss aversion, where a gain-loss neutral individual λ = 1. A loss-averse individual has λ > 1, and a gain-seeking individual has λ < 1 (Supplementary Figure 4).

The probability of selecting the gamble was calculated using the softmax rule:

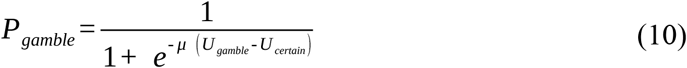

Here, the inverse temperature parameter μ quantifies the degree of randomness in choice behavior. Parameters were fitted for both stimulation ON and OFF conditions, with the following constraints based on our design matrix: *α*_*gain*_ and *α*_*loss*_ ranged from 0 to 2, λ ranged from 0.5 to 3, and μ ranged from 0 to 5.

Trial-by-trial behavioral data (gambles, outcomes, and choices) were preprocessed in Python and passed to Stan as input. For stimulation ON we ran 4 Markov Chain Monte Carlo (MCMC) chains with 2,000 iterations each (1,000 warm-up, 1,000 sampling) using No-U-Turn Sampler (NUTS) for efficient convergence. For stimulation off we adjusted the sampling parameters to ensure robust convergence. We increased the warm-up iterations to 2,000 to allow better adaptation of the sampler, we set the sampling iterations to 4,000 to improve posterior estimation and we adjusted adapted delta to 0.9 to reduce divergences and improve sampling efficiency. This parameter controls the target acceptance rate during the warmup phase of the Hamiltonian Monte Carlo (HMC) sampler. Convergence was assessed using R-hat statistics for all conditions (<1.1), effective sample size (ESS), and posterior distributions were analyzed in Python using ArviZ, and model fits were validated through posterior predictive checks.

### Gain-Loss Learning Model

To determine whether outcomes from previous gamble choices influence subsequent decisions, we fit an alternative gain-loss learning model. This model extends the standard prospect theory framework by including two additional parameters that modulate the subjective value of gambles based on the outcome of the most recent gamble of the same type (gain or loss trial).

Specifically, the utility of the gamble was modified as follows:

In gain trials (CR>0), utility was defined as:

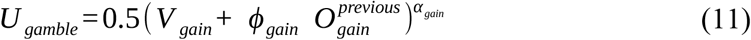

For loss trials (CR<0):

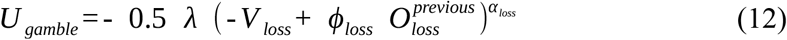

In mixed trials (CR=0), where the certain option is neutral and no history-based modulation is applied, the utility followed standard prospect theory as in (9).

Where *ϕ*_*gain*_ and *ϕ*_*loss*_ are free parameters capturing the influence of previous outcomes. 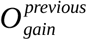 was coded as +1 if the previous gain gamble was a win, and –1 if a loss. Similarly, 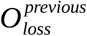 was +1 if the previous loss gamble resulted in a loss being avoided, and –1 if it was incurred.

The certain option was evaluated using the standard prospect theory value transformation as in (9). The probability of choosing the gamble was computed via the softmax rule as in (10).

All parameters were estimated hierarchically using a Bayesian approach. Subject-level parameters were drawn from group-level priors, with bounded constraints imposed via probit-transformed Gaussian priors using Phi_approx(). Inference was performed in Stan using the No-U-Turn Sampler (NUTS). Convergence was assessed using R-hat statistics and effective sample sizes. Posterior predictive checks and trial-level log-likelihoods were used to evaluate model performance. This model allowed us to assess whether past outcomes dynamically biased valuation of new gambles, and whether this was modulated by stimulation (e.g., via dopaminergic effects).

As a control, we ran a third model where *O* ^*previous*^ reflected the outcome of the immediately preceding trial, regardless of whether it was a gamble or a certain choice. This model yielded results qualitatively like the gamble-only version, suggesting that the observed effects are not exclusively tied to previous risk-taking behavior (Supplementary Figure 4).

### Null Learning-Only Model

To isolate the influence of prior outcomes on current decisions without incorporating value-based decision signals from prospect theory, we fit a simplified null model focused solely on learning-related effects from previous gamble outcomes. This model does not include utility computations based on gains or losses but instead examines whether subjects are more likely to gamble following positive or negative feedback on previous trials.

The model assumes that the subjective value of gambling on a given trial is modulated by the outcome of the most recent gain or loss gamble, depending on the trial context. Specifically, the utility of gambling was modeled as a linear function of the previous outcome:

In gain trials (CR>0), utility was defined as:

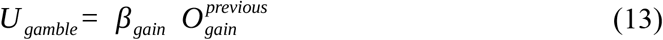

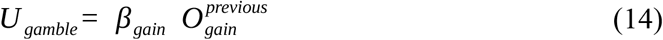

In loss trials (CR<0), utility was:

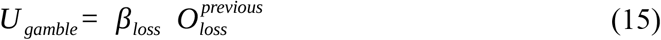

In mix trials (CR=0), utility was set to 0.

Here, 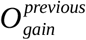 and 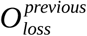 were coded as +1 for positive and –1 for negative outcomes on the most recent gain or loss gamble, respectively. The parameters *β*_*gain*_ and *β*_*loss*_ capture each subject’s tendency to become more or less likely to gamble following a win or a loss in gain or loss trials. Choices were modeled with a softmax function using a subject-specific inverse temperature parameter *μ*, which determines choice stochasticity:

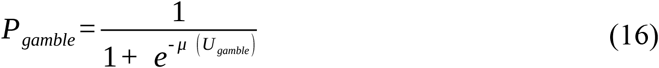

All parameters were estimated using a hierarchical Bayesian approach. Subject-level parameters *β*_*gain*_, *β*_*loss*_ and *μ* were drawn from group-level normal distributions, and *μ* was constrained to be positive via an exponential transformation. The model was implemented in Stan using Hamiltonian Monte Carlo (HMC) sampling via NUTS.

We ran four Markov chains with sufficient warm-up and sampling iterations to ensure convergence, assessed using R-hat values (<1.1) and effective sample sizes.

### Bias-Only Null Model

As a baseline comparison, we implemented a bias-only null model to estimate each subject’s overall propensity to choose the gamble option, independent of task variables such as gains, losses, or prior outcomes. This model assumes that subjects make decisions according to a fixed internal bias toward or against gambling, represented as a latent choice probability on the logit scale.

For each trial, the probability of choosing the gamble was modeled as:

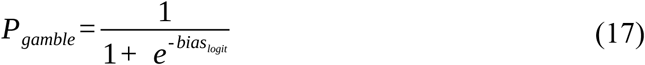

where *bias*_*logit*_ is a subject-specific bias parameter.

These bias parameters were estimated hierarchically. Each subject’s bias was drawn from a group-level Gaussian distribution:

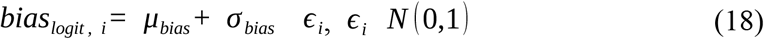

Here *μ*_*bias*_ and *σ*_*bias*_ represent the group mean and standard deviation of bias across subjects, with weakly informative priors:

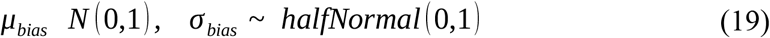

This hierarchical structure accounts for individual variability while borrowing strength across subjects.

Inference was performed in Stan using the No-U-Turn Sampler (NUTS) via CmdStanPy. Convergence diagnostics, including R-hat and effective sample size (ESS), were evaluated to confirm reliable estimation. Model performance was assessed using trial-level log-likelihoods, calculated in the generated quantities block for use in model comparison (e.g., LOO or WAIC).

### Model Comparison

To evaluate and compare the goodness-of-fit of competing computational models, we used Bayesian model comparison based on leave-one-out cross-validation (LOO-CV). We implemented this using the az.compare() function from the ArviZ Python package, which estimates each model’s predictive accuracy while penalizing model complexity.

We compared the five models described in this section for each stimulation condition (ON and OFF): Prospect Theory (standard 4-parameter model), PT Gambling History (Prospect Theory + gain/loss learning history), Null PT Learning (history-only modulation without value terms), Null (bias-only model) and PT Immediate History (alternative variant of the PT Learning model taking the immediate trial).

Model comparisons were conducted on cleaned datasets to ensure high-quality posterior estimates. For both STIM OFF and STIM ON conditions, we computed the expected log predictive density (ELPD) using Pareto-smoothed importance sampling LOO (PSIS-LOO) and combined model weights using Bayesian bootstrap pseudo-Bayesian model averaging (BB-pseudo-BMA).

The method=“BB-pseudo-BMA” option in ArviZ combines model weights based on ELPD values, accounting for uncertainty across models. Lower LOO scores indicate better out-of-sample predictive performance. All comparisons were visualized using ArviZ’s plot_compare() function to aid in interpretation.

### Parameter recovery analysis

To assess the recoverability of model parameters, we conducted a parameter recovery analysis by simulating behavioral data and fitting the model to evaluate how well the estimated parameters matched their true input values (Danwitz et al., 2022).

Two separate datasets for on and off were generated consisting of the same number of trials and patients as the real dataset, 21 patients 50 trials for Prospect theory Model. For the augmented learning Prospect Theory Models we had to increase the number of trials to 150 and patients to 50 in order to get reliable parameter recovery results. Following model fitting, we extracted the hyperprior mean or group-level mean parameter (μ_pr_) and the hyperprior standard deviation or group-level standard deviation (σ_pr_) of the group parameters from the fitted Stan model. This was done by loading individual Markov Chain Monte Carlo (MCMC) samples and computing the mean estimates for alpha gain (*α*_*gain*_), alpha loss (*α*_*loss*_), loss aversion (λ), and decision temperature (τ) from the μ_pr_ variable in Stan and the standard deviations for the same parameters from the σ_pr_ variable in Stan.

Using the group-level posterior distributions, we generated individual-level samples for each parameter. We sampled individual-level values from a normal distribution with mean μ_pr_ and standard deviation σ_pr_ derived from the posterior estimates. We transformed sampled values using a cumulative normal distribution (normcdf) to constrain them within a specified range (e.g., 0 to 2 for alpha gain and alpha loss, 0 to 3 for loss aversion and 0 to 5 for temperature) to apply the same constrains applied in the original model fit to real data. We repeated this process for the model parameters to generate a full set of parameter samples. The sampled parameter values were then used to simulate decision-making behavior using the same prospect theory models that was fitted to the real data. For each simulated subject, choices were generated based on the softmax decision rule incorporating the sampled values of the model parameters. The same model fitted to the patients’ data was fitted to the simulated data and results from both were compared to assess whether the recovered behavioral patterns aligned with those observed in real data.

### Posterior Predictive Check (PPC)

For each participant p we used the original OFF and ON trial sequences exactly as presented (no re-ordering or resampling). Each trial n provides: a certain reward *CR*_*pn*_; two gamble outcomes (*g* 1_*pn*_, *g* 2_*pn*_); a realized-outcome index, when recorded, *outIdx*_*pn*_∈{1,2}; and a condition flag *cond*_*pn*_∈{OFF,ON}. The number of simulated trials per participant and condition matches the observed number. The utilities for the prospect theory model were computed according to either equations (8)-(9) or (11)-(12) depending on the model chosen. And the probability of selecting the gamble was calculated using the softmax rule in (10). We generated D=500 posterior-predictive replicates per participant (default). When full posterior samples were available, each replicate drew one posterior sample for that participant (for both conditions) and used it to simulate all of the participant’s trials in that replicate. When only point estimates were available, the same estimates were reused across replicates. For each replicate, OFF trials were simulated with OFF parameters and ON trials with ON parameters. If the simulated choice selected the certain option, the payoff was *CR*_*pn*_. If it selected the gamble, we used the *outIdx*_*pn*_ to pick *g* 1_*pn*_ or *g* 2_*pn*_. Across the D replicates, we summarized for each participant the median EV−CR|accepted (OFF, ON), and the median earnings per trial (OFF, ON). Medians across replicates reduce the influence of rare extreme gamble outcomes. For the group level statistics, we report paired *t*-test on ON–OFF differences, Wilcoxon signed-rank test, Cohen’s d_z_ for paired designs, 95% bootstrap confidence interval for the mean difference (5,000 resamples).

### Decomposition of ON–OFF changes in loss-side slope

We asked whether stimulation primarily alters loss aversion (λ) or the loss-domain curvature parameter (*α*_*loss*_, abbreviated α). On the loss side (x<0), the Prospect Theory value function is U(x) = −λ (−x) α. The (absolute) slope at loss magnitude L=|x| is:

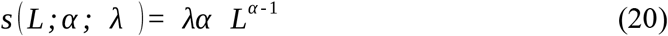

For numerical stability we analyze the log-slope,

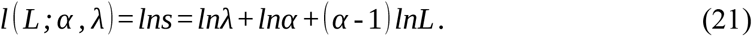

We evaluate slopes at a fixed loss x0=−20, so L=|x0|=20 and *L*_*x*_ = ln *L* = ln 20. For each participant *i* with OFF parameters 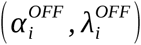 and ON parameters 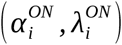, the ON–OFF change in log-slope decomposes exactly as:

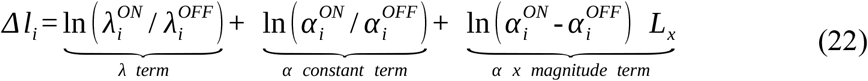

We summarize the α contribution as the sum of its two pieces (second and third terms). To assess which component “explains” the total change Δ*ℓ*, we report (i) Pearson correlations between Δ*ℓ* and each component (λ term; α contribution), and (ii) an attribution regression,

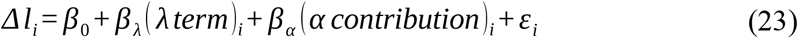

with standardized coefficients and partial R^2^. Group inference uses two-tailed one-sample *t*-tests on each component (mean vs 0) with 95% CIs by bootstrap (Figure 4d-e).

### Counterfactual Parameter Decomposition analysis

To identify which model parameter(s) most plausibly mediate the stimulation effect, we computed, on each participant’s ON trials, the expected behavior under hypothetical “bundles” of parameters in which exactly one parameter is switched from its OFF posterior to its ON posterior while all other parameters are held at their OFF posteriors. We quantified the change in two decision metrics relative to an all-OFF baseline: (i) expected EV−CR among accepted gambles and (ii) expected payoff per trial. We repeated the analysis separately for gain-context trials (CR>0) and loss-context trials (CR<0). For participant p and trial n (restricted to that participant’s ON-condition trials) we have certain reward *CR*_*pn*_; two gamble outcomes (*g* 1_*pn*_, *g* 2_*pn*_); task expected value Task expected value *EV*_*pn*_ = 0.5 *g* 1_*pn*_ + 0.5 *g* 2_*pn*_, advantage *EV*_*pn*_ - *CR*_*pn*_. Posterior samples (draws) over parameters indexed by s=1,…,S. Let *G*_*p*_ = *n* : *CR*_*pn*_ > 0 be the participant’s ON trials in the gain context, and *L*_*p*_ = *n* : *CR*_*pn*_ < 0 the loss context. For each draw s, the utilities for the prospect theory model were computed according to either equations (8)-(9) or (11)-(12) depending on the model chosen. And the probability of selecting the gamble was calculated using the softmax rule in (10). Then we use the posterior-mean gamble probability on each trial:

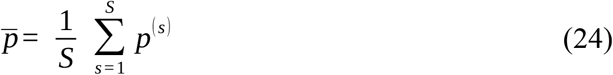

A parameter bundle selects, for each parameter, whether to use the participant’s OFF posterior or ON posterior. For the no-history model the parameter set is {αgain, αloss, λ, τ}, and for the history model {λ, αgain, αloss, ϕgain, ϕloss, τ}. The baseline bundle (all-OFF) evaluates each ON trial using that participant’s OFF posteriors for all parameters to obtain 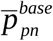, and flip-j bundle was computed switching only parameter j to the ON posterior while keeping all other parameters at OFF, yielding 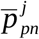. This isolates the unique contribution of parameter. We define a set of trials *M* ∈ *G*_*p*_, *L*_*p*_and we computed the expected EV−CR among accepted gambles (probability-weighted) as:

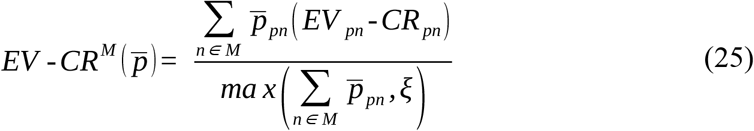

And expected payoff per trial as:

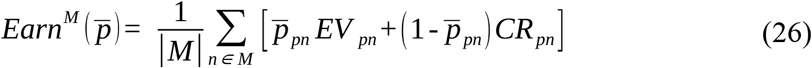

Where *ξ* is a small constant to guard against division by zero when no gambles are predicted. We compute these metrics twice per context: once under the baseline probabilities 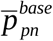 and once under the flip-j probabilities 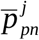.

We computed the subject level counterfactual effects by defining for each participant p, parameter j and context *M* ∈ {*G*_*p*_, *L*_*p*_}:

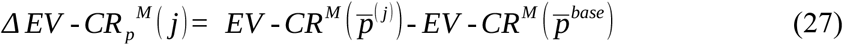

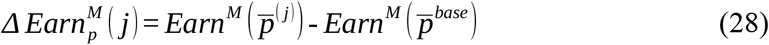

These deltas populate a subjects × parameters matrix for each metric and context (gain, loss). Across participants, we summarize Δ values with the mean ± s.e.m., 95% CI, and a two-tailed one-sample t-test of Δ=0 for each parameter j within each context. Bar-and-whisker plots (means ± 1.96·s.e.m.) of Δ facilitate visual comparison of parameters.

### Computation of subjective utility vs Objective outcome: Value function

For each participant p and condition c ∈ {OFF,ON}, we assembled trial-wise pairs 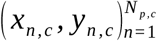 where *x*_*n, c*_ is the objective outcome (task units) and *y*_*n, c*_ is an empirical utility estimate for that trial (as computed in the main modeling pipeline). Trials with non-finite x or x were excluded. Fits were attempted only when ≥6 valid points were available in a condition.

The prospect theory function used for each condition c, subjective value was modeled as a piecewise power function with loss aversion:

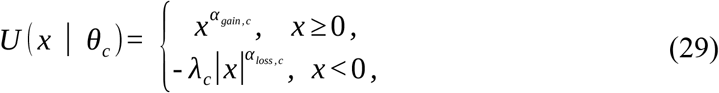

with parameter vector *θ*_*c*_ = (α_gain,c_, α_loss,c_, λ_c_). Parameters were estimated by bounded nonlinear least squares,

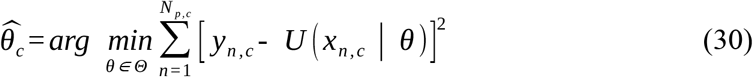

with bounds α_gain_, α_loss_∈[0,2] and λ∈[10^−3,10^]. The initial guess was (0.9, 0.9, 2.0). We used MATLAB’s lsqcurvefit (R2018) with MaxIterations=1000 and MaxFunctionEvaluations=5000. For visualization of an example participant, we evaluated the fitted curves on a dense grid x ∈ [−20,20] and overlaid the scatter of (x, y).

To summarize shape differences between conditions at the group level, we computed subject-level utility curves on a common grid x ∈ [−20,20] using each participant’s fitted parameters:

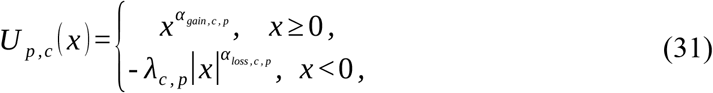

We then averaged utilities across participants at each xxx and formed pointwise 95% confidence bands by nonparametric bootstrap over participants (resampling subjects with replacement, B=2000):

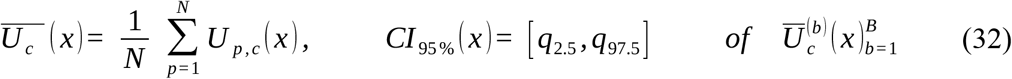

Optional “spaghetti” lines (thin individual curves) were drawn for transparency; shaded ribbons show the bootstrap CIs; vertical and horizontal reference lines mark x=0 and U=0.

### Probability of gamble observed vs Utility difference on an OFF-baseline axis

For each participant p with trials n=1…N_p_ we first computed, under that participant’s OFF parameters only independent of whether the actual trial was ON or OFF, the trial-wise subjective-utility difference using equations (8)-(10). That OFF-parameter ΔU is the common x-axis.

Observed choices were coded y_pn_=1 for gamble and 0 for certain. We then fit a logistic regression of observed choice on the OFF-axis utility difference,

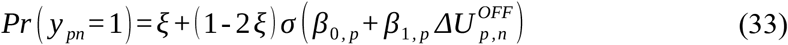

where σ(·) is the logistic function and ε=0.02 is a symmetric lapse (fixed). The on the OFF axis is the ΔU value at which the fitted curve crosses 0.5:

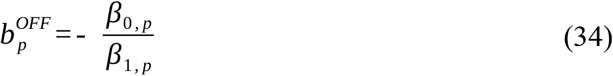

To isolate the impact of the stimulation-related change in loss-curvature, we built a counterfactual probability sequence for the same trials by flipping only α_loss_ from OFF to each participant’s ON estimate, while keeping α_gain_, λ, and the softmax slope τ at their OFF values:

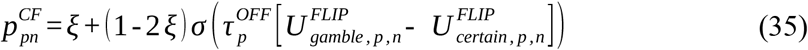

With *U* ^*FLIP*^ identical to the OFF formulas but replacing only 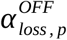 by 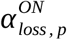. For visualization on the same OFF ΔU axis, we summarized this counterfactual with a logistic-in-ΔU^OFF^,

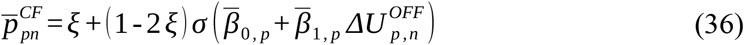

And plotted the correspondence boundary:

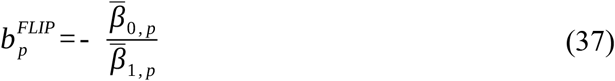

This produces two directly comparable curves—observed OFF and counterfactual αloss→ON— on a common horizontal axis ΔU^OFF^, making the horizontal boundary shift interpretable as the isolated effect of increasing loss-side curvature. Plotting used a data-driven ΔU^OFF^ grid spanning the [1,99]th pooled quantiles with light padding.

For group summaries, we first defined a common ΔU^OFF^ grid from the pooled OFF ΔU values across participants (quantiles as above). For each participant p, we evaluated the fitted OFF curve 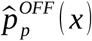 and the fitted counterfactual curve 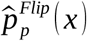 along that grid. We then computed mean **±** SEM across participants at each grid point:

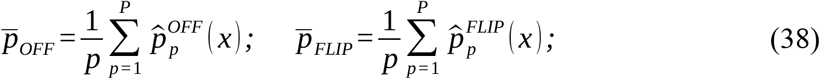

with SEM from the across-participant standard error. Participant-specific boundaries 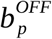 and 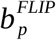 were summarized by their means (and can be inferentially compared via 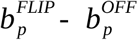).

### Schematic figures (Prospect Theory value and decision mapping)

For the model used in all schematic panels we used the same Prospect Theory value function specified earlier (29) to compute subjective utility for a single outcome x. For a 50–50 gamble with outcomes (g1, g2) and a certain option cr, the utilities are

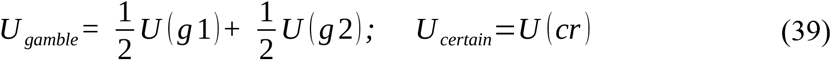

and the decision variable is the utility difference ΔU = *U*_*gamble*_ - *U*_*certain*_,

And choice probability was computed as in (9). To anchor the diagrams, we used one representative trial with c=−5, g1=0, g2=−15, and we marked (g1, g2, c) on the value curve.

For the single condition schematic in Figure 3a, the panel illustrates the shape of the group-typical value function and its mapping to choice without contrasting conditions. To illustrate the parameters, we instantiated the curve with the sample-average OFF parameters (mean across participants p) α_gain_, α_loss_ and λ. The softmax slope used for the central decision curve equals the sample-average μ; two faint companion curves at μ_low_=0.45 and μ_high_=1.80 illustrate the effect of slope on P(gamble).

In the top figure u(x) over x ∈ [−20,20] with vertical/horizontal zero guides, and with the three stimulus values (g2, c, g1) projected onto the curve. In the bottom figure we plot the decision process as P(gamble) over ΔU ∈ [−12,12], with the same μ using formula (10). Note that this figure is didactic (not an inferential test): it visualizes the group-typical concavity for gains, convexity for losses weighted by λ, and the monotonic mapping from ΔU to choice.

In Figure 4a OFF vs ON schematic (two-condition comparison) we show how increasing loss-side curvature in ON shifts both the value curve and the decision mapping when evaluated on a common OFF scale. We held α_gain_, λ, μ fixed across conditions at their OFF group means. ON differed only in α_loss_ being multiplied a factor of f = 1.90. In the top (value curves), we plotted u_OFF_(x) and u_ON_(x) on x ∈ [−20,20]. Because α_loss_ON>α_loss_OFF, the ON loss branch is steeper (more loss-sensitive), while the gain branch (same α_gain_ is unchanged. The three stimulus values (g2, c, g1) are marked on both curves.

In the bottom of Figure 5a we plot Bottom P(gamble) on a common OFF axis. To place both conditions on a single scale, we used a shared ΔU_OFF_. For each participant p, we swept a grid of certain values c ∈ [−30,5] while holding the same representative gamble (g1, g2) = (0, −15). For each grid point, we computed ΔU_OFF_ and the corresponding ON counterfactual where ON replaces only α_loss_ for each patient by f·α_loss_; all other parameters use each participant’s OFF estimates. We then mapped both ΔU series to probabilities with the participant’s OFF slope μ using (10). Finally, we averaged P(p) across participants at each x-value. The result is a pair of mean curves P_OFF_(ΔU_OFF_) and P_ON_(ΔU_OFF_) shown on the shared OFF axis. With only α_loss_ increased, ON shifts the value of loss-side outcomes downward relative to OFF; on the common OFF axis this moves the ON decision curve leftward (higher gamble probability at the same ΔU_OFF_ magnitude for losses).

## Acknowledgments

We gratefully acknowledge the willingness of our patients to participate in this study. We thank the Cedars-Sinai operating room staff for their assistance, in particular Robert Zelaya. We thank John O’Doherty and Rob Rutledge for insightful discussions and comments on the experimental design and interpretation of the results, and Michael Kyzar and Wenying Zhu for assistance with data acquisition and processing. This work was funded by the NIH (U01NS132788 and U01NS098961 to U.R. and A.N.M.), Cedars-Sinai Department of Neurosurgery bridge funds, and the Simons Foundation SCGB.

## Author Contributions

Conceptualization, M.Y., C.M., U.R.; Methodology, M.Y., Z.F., U.R.; Data acquisition, M.Y., C.M., H.C., A.M.; Data curation, M.Y.; Formal analysis, M.Y.; Software, M.Y.; Visualization, M.Y.; Writing—original draft, M.Y., U.R.; Writing—review & editing, A.M., Z.F., M.T., U.R.; Supervision, U.R.; Clinical oversight, M.T., A.M.; Electrode localization, C.M., S.C.; Project administration, M.Y., U.R. Funding acquisition, U.R,

## Supplementary Figures

**Supplementary Figure 1.**
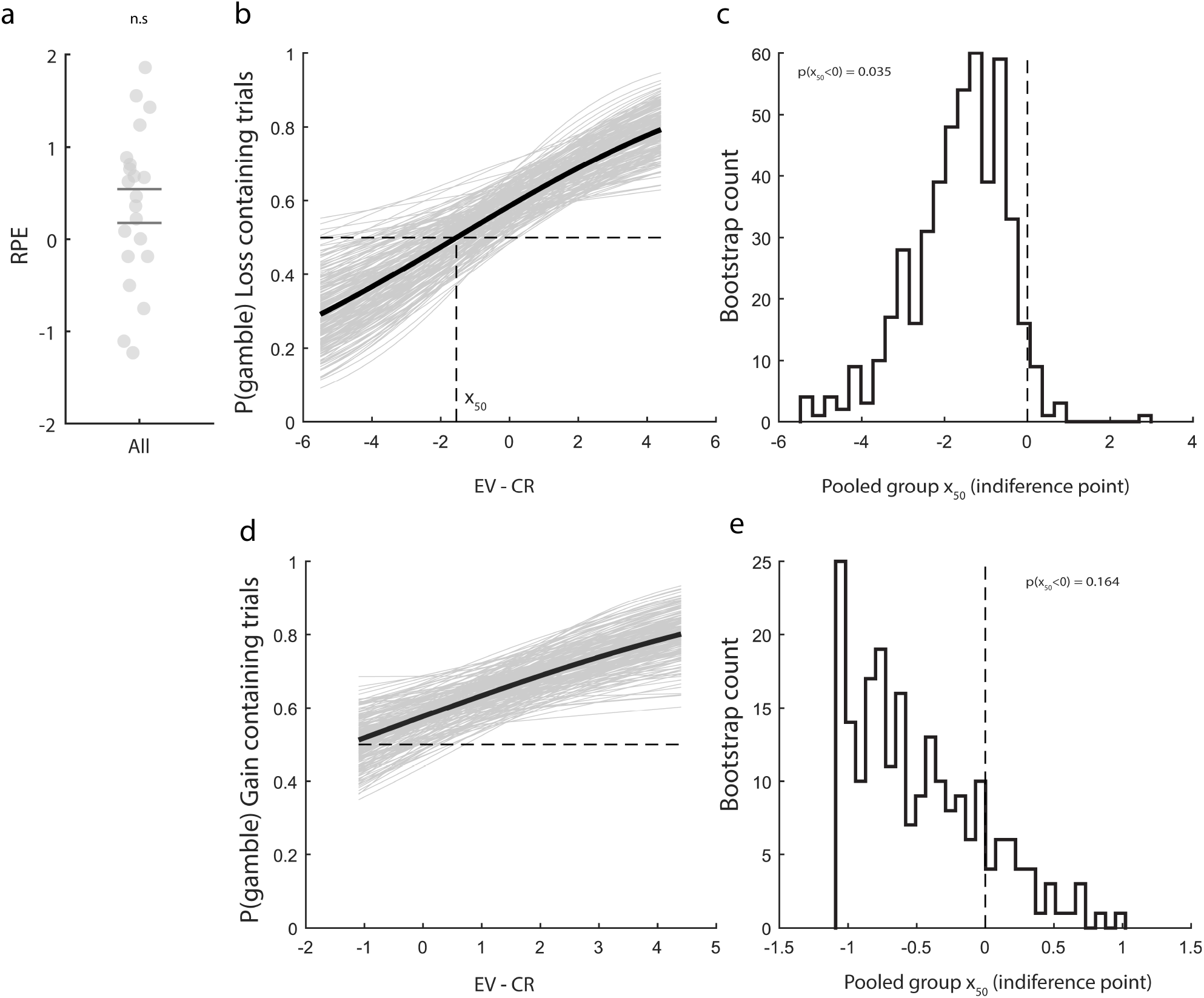
Reward prediction errors and domain-specific choice behavior. (a) RPE (reward prediction error = Outcome − EV), pooled across trials, centered around zero as expected. Asterisks denote two-tailed paired t-test significance (*p < 0.05, *p < 0.01; “n.s.” = not significant). (b) Psychometric function for loss-containing trials (mixed and pure-loss), collapsing across ON and OFF conditions. Gray lines indicate bootstrap resamples; black line indicates the group-average fit from a logistic model with subject intercepts. The horizontal dashed line indicates indifference (P(gamble) = 0.5), and the vertical dashed line indicates the indifference point (x_50_). A negative x_50_ reflects loss-seeking behavior in the loss domain. (c) Bootstrap distribution of the pooled indifference point (x_50_) for loss-containing trials. The dashed line indicates x_50_ = 0. The distribution is shifted below zero, consistent with loss-domain risk-seeking behavior (one-sided bootstrap p = 0.035). (d) Psychometric function for gain-containing trials (mixed and pure-gain), collapsing across ON and OFF conditions. Gray lines indicate bootstrap resamples; black line indicates the group-average fit from a logistic model with subject intercepts. The horizontal dashed line indicates indifference (P(gamble) = 0.5). The curve does not cross the indifference line within the observed ΔEV range, indicating no well-defined indifference point. (e) Bootstrap distribution of the pooled indifference point (x_50_) for gain-containing trials. The dashed line indicates x_50_ = 0. The distribution is not reliably shifted from zero, consistent with the absence of a systematic bias in the gain domain (one-sided bootstrap p = 0.164).

**Supplementary Figure 2.**
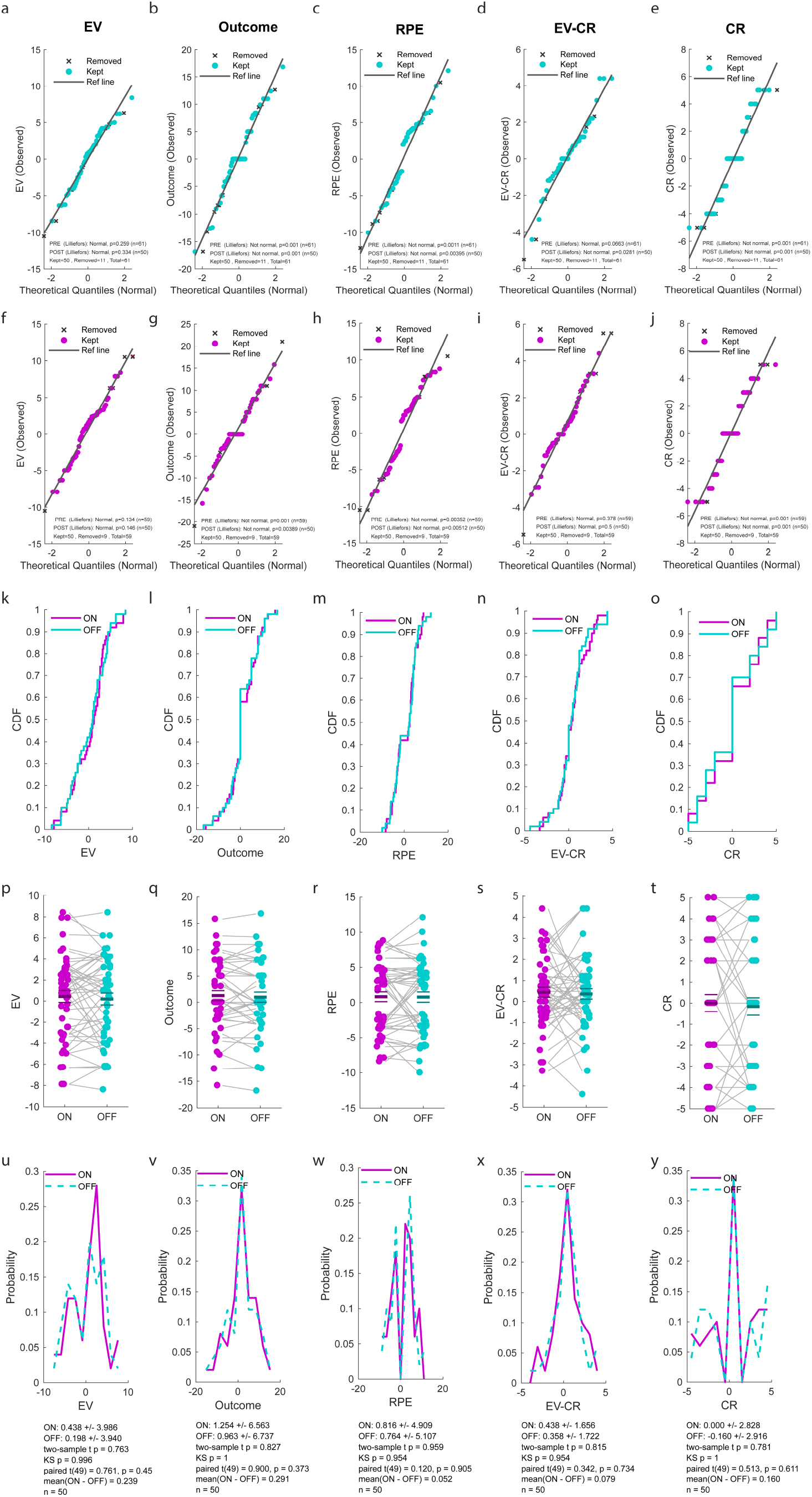
Control analyses for a balanced experimental design between stimulation ON and OFF trials. **(a –e)** QQ-plots for OFF trials showing the distribution of EV, Outcome, RPE = Outcome − EV, EV − CR (objective value advantage), and CR. Dark gray “×” marks indicate trials removed by the balancing step; filled symbols indicate kept trials; the black line is the OLS reference. Panel text reports normality tests and sample sizes (PRE = before balancing, POST = after). **(f–j)** QQ-plots for ON trials, same variables and conventions as in (a–e). **(k–o)** Empirical CDF overlays for ON (magenta) and OFF (cyan) after balancing. The near overlap of the curves indicates that the stimulus distributions are matched across conditions for all five variables. **(p–t)** Participant-level paired summaries (one dot per participant; gray lines link ON and OFF means within participant) for EV, Outcome, RPE, EV−CR, and CR. Horizontal ticks show group means. These panels verify no systematic ON –OFF differences in the stimulus features presented. **(u–y)** Histogram outlines showing the distribution of EV, Outcome, RPE = Outcome − EV, EV − CR (objective value advantage), and CR (ON = solid magenta, OFF = dashed cyan) with summary statistics and tests printed in each panel (two-sample t and KS; paired t on participant means where relevant). Together, these checks demonstrate that the experimental inputs were balanced between ON and OFF sessions—i.e., participants saw comparable EV, outcomes, RPE, EV−CR, and CR in both conditions—so any ON–OFF differences in behavior cannot be attributed to differences in the stimulus distributions. Color code used throughout: OFF =cyan; ON = magenta.

**Supplementary Figure 3.**
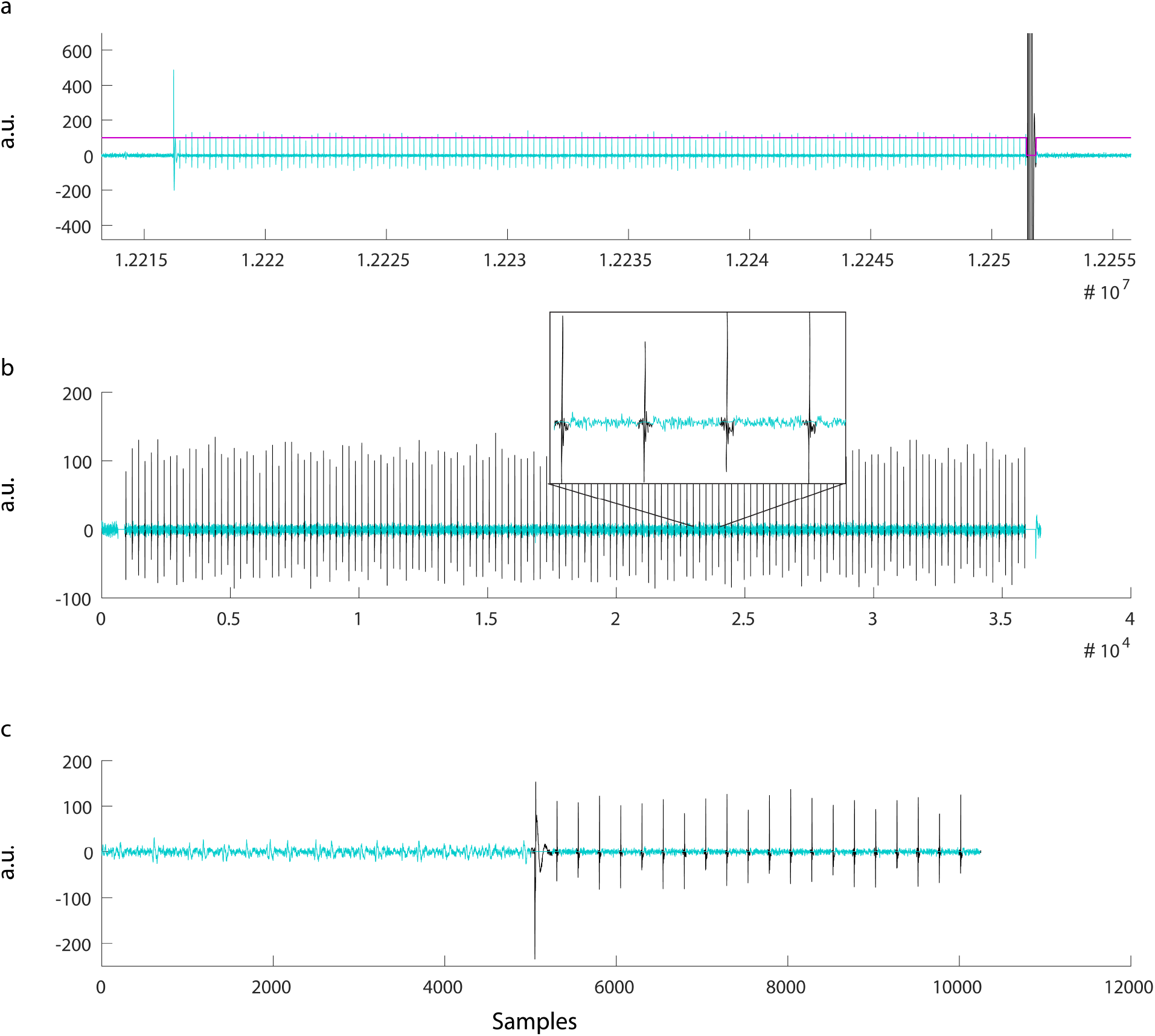
Stimulation artefact removal. **(a)** Example trace of a stimulation train showing the zero padded sniped signal for the last big artefact of the stimulation train artefact at the end in the first step of the artefact removal algorithm. Raw filtered signal in black, cleaned signal in blue clipping mask in magenta. **(b)** Snipped signal for the second step of the stimulation artefact removal algorithm. Raw filtered signal in black, cleaned signal in blue. Inset: zoomed signal. **(c)** Example trace for the third step of the stimulation artefact removal algorithm showing the removal of a longer portion of the artifact for the bigger pulses found at the beginning of the stimulation train pulses. In black the original signal is shown, in blue the cleaned signal.

**Supplementary Figure 4.**
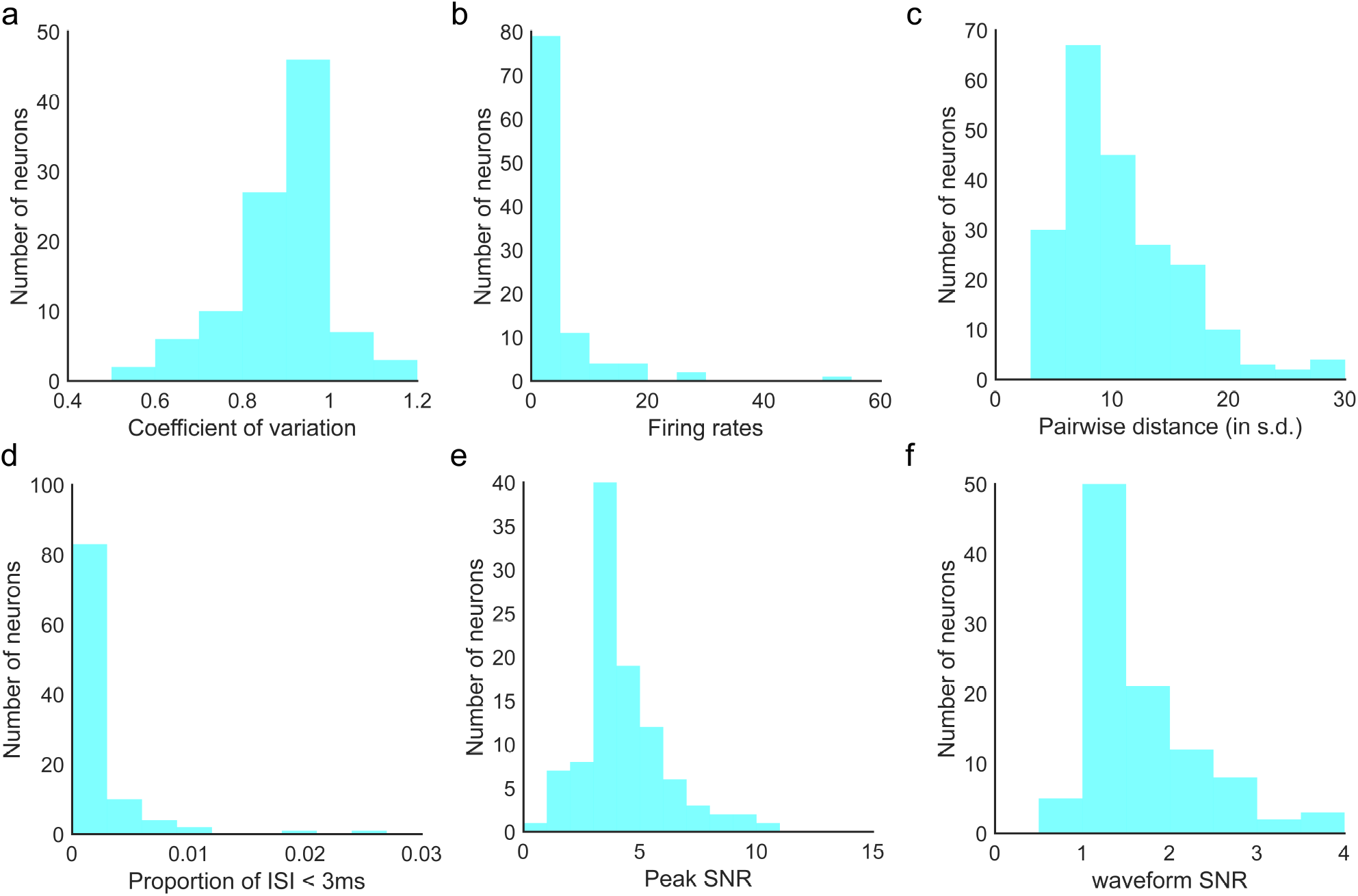
Spike sorting quality metrics for all identified putative single cells. **(a)** Coefficient-of-variation in the ISI for each identified putative single cell (0.90± 0.12, mean ± s.d.). **(b)** Average firing rate within the entire recording session for all identified putative single cells (4.17± 7.44, mean ± s.d.). **(c)** Pairwise isolation distance between putative single cells identified from the same wire (10.77± 5.14, mean ± s.d.). **(d)** Proportion of inter-spike intervals (ISI) that were shorter than 3ms (0.002± 0.004, mean ± s.d.). **(e)** Waveform peak signal-to-noise ratio (SNR), which is the ratio between the peak amplitude of the mean waveform and the s.d. of the noise of each identified putative single cell (4.28± 1.80, mean ± s.d.). **(f)** Waveform signal-to-noise ratio (SNR), which is the ratio between the amplitude of the mean waveform and the s.d. of the noise of each identified putative single cell (1.70± 0.65, mean ± s.d.).

**Supplementary Figure 5.**
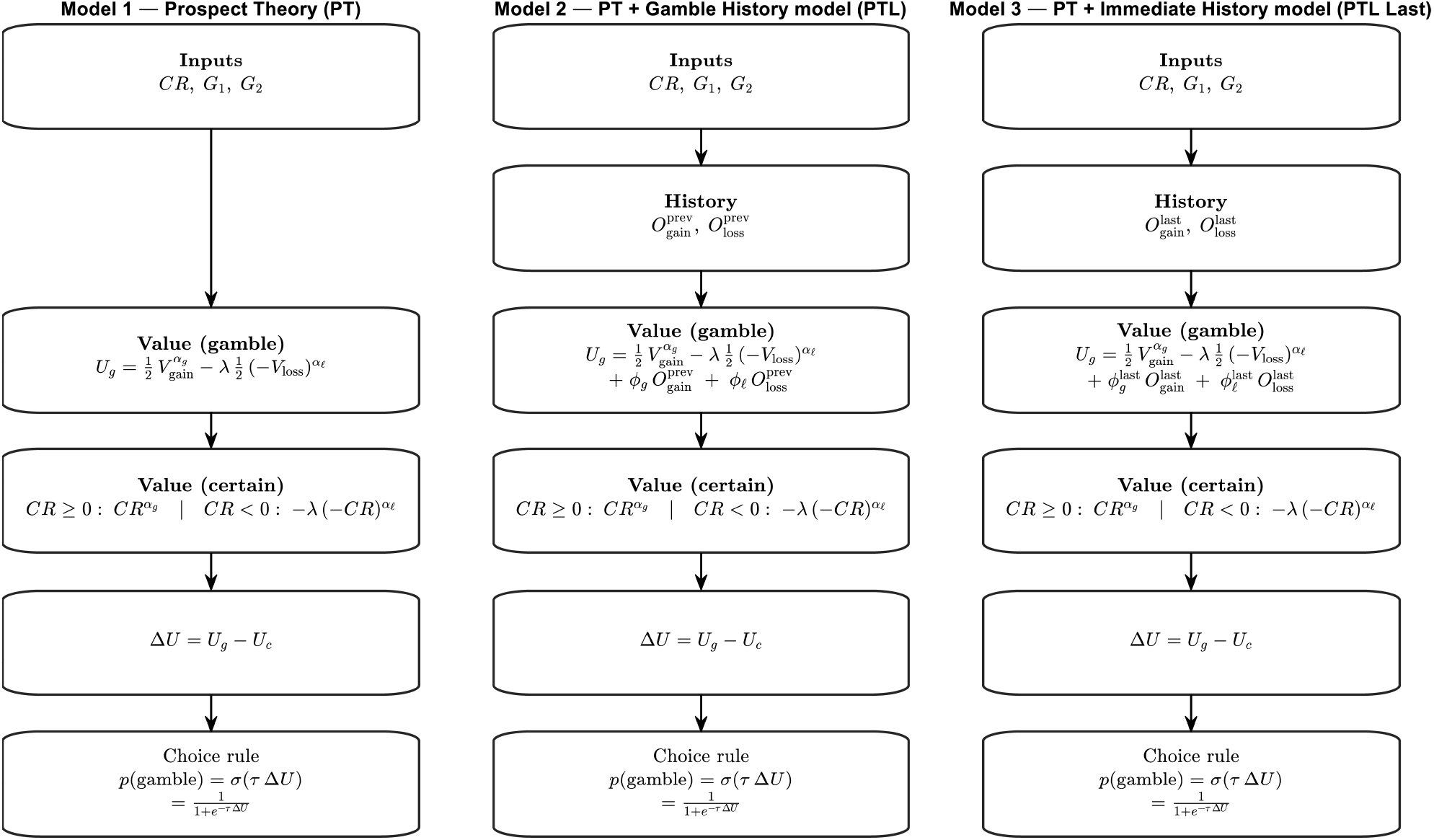
Prospect-theory models with outcome-history terms. Each column shows the computation from inputs to choice probability. Model 1 (PT) uses a standard prospect-theory value function for a 50/50 gamble with two outcomes G_1_, G_2_. Choice depends on the value difference ΔU via a logistic rule. Model 2 (PT + Gamble History) augments the gamble value with linear carry-over of the previous trial’s realized outcome in the gain and loss domains. Model 3 (PT + Immediate History) is identical but uses the immediate last trial only. Notation. G1 is the offered gamble in the upper right corner on the screen, G2 G1 is the offered gamble in the lower right corner on the screen, CR is the certain reward; α _g_, α_*ℓ*_ are curvature parameters (gains, losses), λ is loss aversion, τ is inverse temperature. 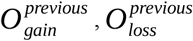 denote the realized outcome on the referenced trial, coded by domain (magnitude in that domain, zero otherwise). Positive ϕ parameters capture attractive recency (recent gains increase; recent losses decrease); negative ϕ capture repulsive recency.

**Supplementary Figure 6.**
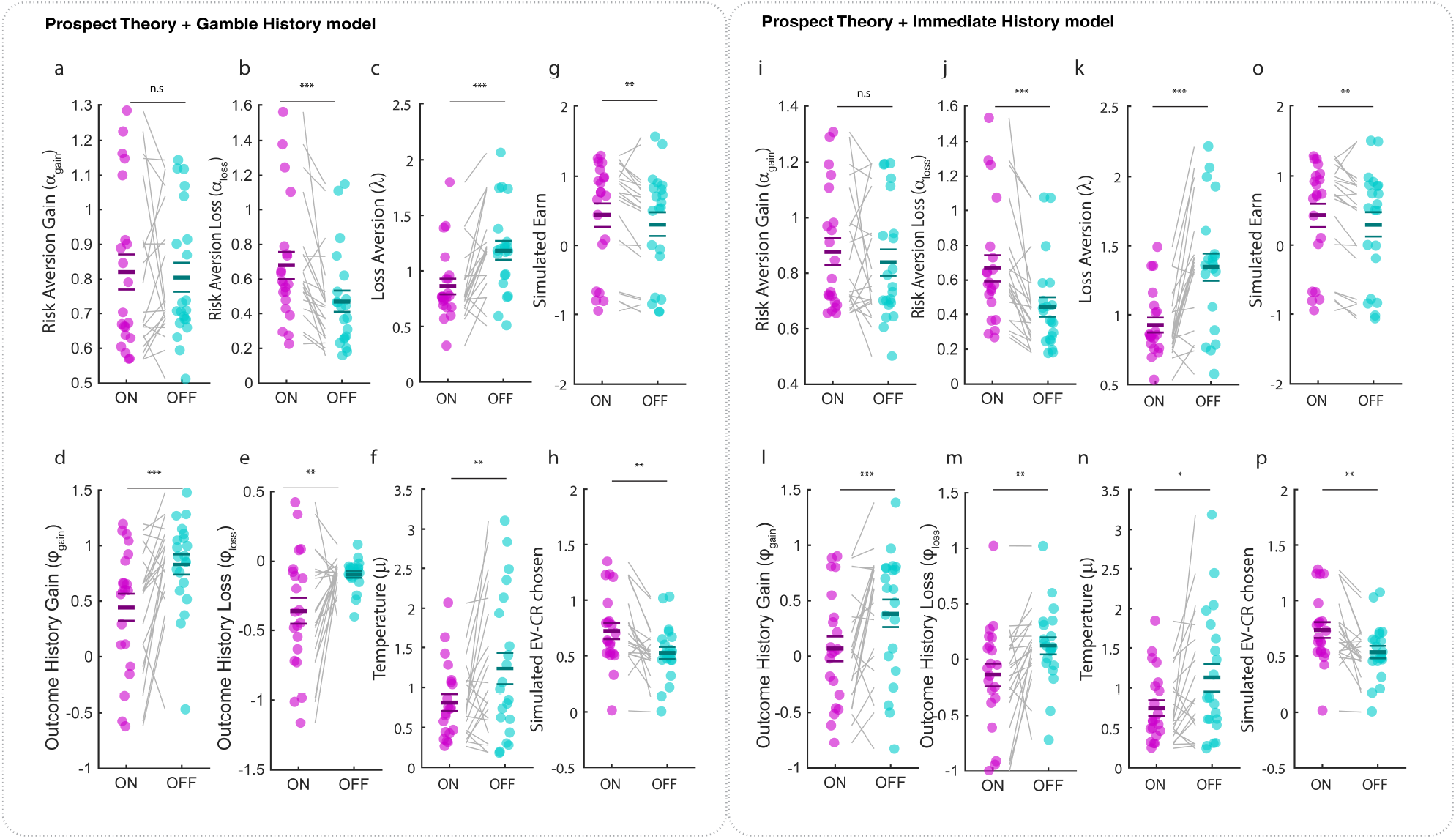
Prospect Theory models with history terms: parameter comparisons under stimulation. Left block — Prospect Theory + Gamble-History model. The standard Prospect Theory parameters are estimated together with history weights that capture how past outcomes bias current choice (separate weights for gains and losses aggregated over recent trials). Panels show, from left to right and top to bottom: risk aversion in gains αgain, risk aversion in losses αloss, loss aversion λ, simulated earnings, history-gain weight (past gains), history-loss weight (past losses), temperature μ, and simulated EV−CR of the chosen option. Right block — Prospect Theory + Immediate-History model. As above, but the history terms capture only the previous trial (separate last-trial gain and last-trial loss weights). Panels are ordered analogously. In all panels, ON (magenta) and OFF (cyan) show participant-level estimates; gray lines connect the same participant across conditions; horizontal ticks denote means (± s.e.m.). Asterisks indicate two-tailed paired t-tests between ON and OFF (p < 0.05; *p < 0.01; **p < 0.001; n.s., not significant). “Simulated” metrics are posterior-predictive summaries generated from the fitted parameters on each participant’s actual trial sequence. Color code: stimulation OFF cyan; stimulation ON magenta.

**Supplementary Figure 7.**
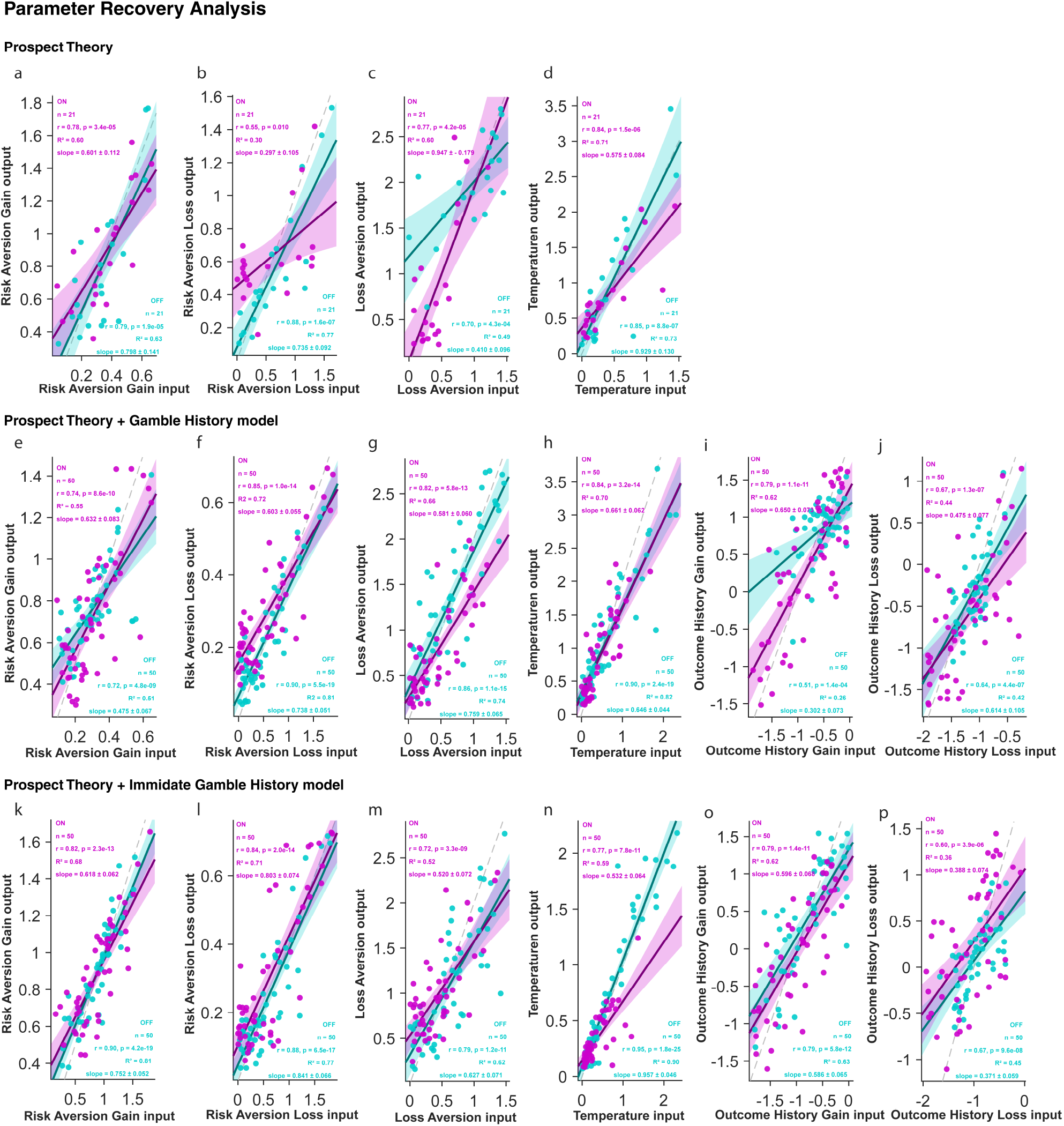
Parameter recovery for Prospect Theory models (with and without history terms). What is shown. Each panel plots recovered (fit) parameters on the y-axis against the ground-truth “input” parameters used to simulate data on the x-axis. Points are per participant/condition simulations; OFF is cyan and ON is magenta. The dashed gray line is the identity (perfect recovery). Solid lines are linear fits with 95% CI bands. For each participant and condition, we simulated choices from the model using the participant’s fitted parameters on their actual trial structure (gains, losses, certain rewards). We then re-fit the same model to the synthetic choices with identical priors/procedure and compared recovered vs. input values. Prospect Theory (top row, a–d): **(a)** risk aversion in gains αgain; **(b)** risk aversion in losses αloss; **(c)** loss aversion λ; **(d)** softmax temperature μ. PT + Gamble-History model (middle row, e–j): **(e)** αgain; **(f)** αloss; **(g)** λ; **(h)** μ; **(i)** history-gain weight; **(j)** history-loss weight (weights capturing how past outcomes bias current choice). PT + Immediate-History model (bottom row, k–p): **(k)** αgain; **(l)** αloss; **(m)** λ; **(n)** μ; **(o)** last-trial gain weight; **(p)** last-trial loss weight. Core Prospect Theory parameters (αgain, αloss, λ, μ) show strong, near-unity input–output relations, indicating they are well identified under our task and priors. History weights are recoverable but noisier (greater scatter, modest shrinkage toward the mean), which is expected given fewer effective samples per weight. Overall, the recovery analyses support that the ON–OFF parameter differences reported in the main text reflect identifiable changes in the underlying constructs rather than fitting artifacts. Color code: stimulation OFF cyan; stimulation ON magenta.

## Supplementary Information

As intended by the task design, reward-prediction errors (RPEs) for gambles were centered near zero at the group level (Figure 2j). A one-sample t-test on the subject means did not differ from zero (*t*_20_ = 2.00; p = 0.06), indicating that outcome surprises were balanced overall and unlikely to explain the ON–OFF earning difference.

### Loss aversion and risk aversion in the learning augmented models

In both versions of the learning augmented models, the stimulation-driven reduction in loss aversion was again observed. In the learning model based on previous gamble outcomes λ was again significantly greater than 1 during OFF (one-sample tests vs 1: *t*_20_ = 2.12 ; p = 0.05), but not during ON (one-sample tests vs 1: *t*_20_ = 1.9574 ; p = 0.06), with a strong ON/OFF contrast (paired t-test: *t*_20_ = -4.68; p = 0.0001) (Supplementary Figure 4c). A similar pattern held in the control model using the immediately preceding trial’s outcome: λ was again significantly greater than 1 during OFF (one-sample tests vs 1: *t*_20_ = 3.54; p = 0.002), but not during ON (one-sample tests vs 1: *t*_20_ = -1.38; p = 0.18403), with a strong ON/OFF contrast (paired t-test: *t*_20_ = -5.40; p = 2.7637 10^-59^) (Supplementary Figure 4k).

### Model comparison and parameter recovery analyses

We compared five models (see Methods) using Leave-One-Out cross-validation computed efficiently with Pareto-Smoothed Importance Sampling (PSIS-LOO) expected log predictive density (ELPD-LOO; larger is better) (Aki Vehtari et al., 2017; A Vehtari et al., 2017). Across ON and OFF sessions, all Prospect-Theory (PT) variants performed similarly, whereas both null models were clearly worse. For the OFF models, the best ELPD was the “PTL-Immediate History” variant (ELPD = −507.62; SE = 13.39). Differences among PT models were small relative to their uncertainty (PT+Gamble History: ΔELPD = 0.92 with SE ≈ 12.75; PT: ΔELPD = 10.26 with SE ≈ 13.48), indicating no reliable separation among PT variants. In contrast, the null models were decisively worse (Null: ΔELPD = 27.83 with SE ≈ 10.31; Null-PTL: ΔELPD = 79.85 with SE ≈ 5.16). Model weights likewise split across the PT family (PT+Immediate History 0.56; PT+Gamble History 0.42; PT 0.02) and were near zero for nulls (≤0.01).

For the stimulation ON models, the “PTL” variant was best (ELPD = −522.05; SE = 12.77). The other PT models were statistically indistinguishable from it (PT: ΔELPD = 1.52 with SE ≈ 12.51; PT+Immediate History: ΔELPD = 3.44 with SE ≈ 12.55). Again, null models were much worse (Null: ΔELPD = 27.33 with SE ≈ 9.47; Null-PTL: ΔELPD = 58.41 with SE ≈ 5.65). Weights were distributed across the PT family (PT+Gamble History 0.55; PT 0.34; PT+Immediate History 0.11) and ∼0 for nulls (≤0.01).

ELPD-LOO model comparison showed that the three Prospect-Theory models were not reliably different from one another (all |ΔELPD| ≤ 10.3 with SEs ≈ 12–13) in either ON or OFF sessions, but all outperformed the corresponding null models by large margins (e.g., OFF Null ΔELPD = 27.8 with SE ≈ 10.3; ON Null ΔELPD = 27.3 with SE ≈ 9.5). We therefore base inference on the Prospect-Theory family and treat the PT variants as comparable in predictive accuracy.

(Note that the rule of thumb used: a model is “reliably worse” than the best if |ΔELPD| > 2×SE. Your ΔELPD values above were computed as ELPD_best_ − ELPD_model_).

As an additional control, we performed parameter recovery from simulated data. For each participant and condition, we simulated choices from the fitted Prospect-Theory models on every trial. On each trial we computed ΔU from the value function, converted it to a choice probability, and averaged across posterior draws. Two Posterior Predictive Check (PPC) summary metrics were then formed per participant: (1) probability-weighted EV–CR among accepted gambles, and (2) expected payoff per trial. We summarized stimulation effects as within-participant ON–OFF differences (Δ) with paired *t*-tests (see Methods).

In the observed data (no simulation), participants accepted gambles with larger EV–CR during ON than OFF and earned more per trial when stimulated Figure 2e,h. These empirical patterns were reproduced by Posterior Predictive Check analyses (PPCs) from the base Prospect Theory Model: EV–CR|accepted increased with stimulation (paired t-test: *t*_20_=3.39, p=0.003) and expected earnings also rose (paired t-test: *t*_20_=2.20, p=0.04).

The history-augmented models yielded the same qualitative conclusions. In Prospect Theory + Gamble History, stimulation increased EV–CR|accepted (paired t-test: *t*_20_=3.68, p=0.001) and increased expected earnings (paired t-test: *t*_20_=2.78, p=0.01). In Prospect Theory + Immediate History model, EV–CR|accepted again increased (paired t-test: *t*_20_=3.68, p=0.001) and expected earnings were higher ON than OFF (paired t-test: *t*_20_=2.87, p=0.01).

Together, the observed data and PPCs converge on the same conclusion: SN stimulation shifts choices toward higher-EV gambles and yields higher payoffs, with medium-to-large within-participant effects that are robust across all three model specifications.

To verify identifiability, we ran a standard parameter-recovery procedure (see Methods). Briefly, for each model and condition (ON/OFF) we (i) drew ground-truth parameters from the empirical posterior, (ii) simulated choices on the original trial sequences, (iii) re-fit the model to those synthetic choices, and (iv) correlated the recovered estimates with the generating values across synthetic participants. For each parameter we report the Pearson correlation r and its p-value (linear fit slopes and R^2^ are shown on the figure; slopes near 1 indicate calibration, high r indicates rank-order recoverability). Overall, recovery was good-to-excellent across models. The history coefficients were particularly well recovered, core Prospect-Theory parameters showed strong recovery for ϕ_loss_, ϕ_gain_, λ, α_gain_ and α_loss_, and temperature μ was also robust. (Table 2.)

**Table 1.** Model comparison using leave-one-out cross-validation (LOO) for the STIM OFF condition. Reported metrics include the expected log pointwise predictive density (ELPD_loo_), the effective number of parameters (p _loo_), the difference in ELPD relative to the best model (ΔELPD), the standard error of the ELPD difference (SE), and the model weight reflecting the posterior probability of being the best model. The Prospect Theory + Gambling History model achieved the highest ELPD and model weight, though the top three models were not statistically distinguishable given their ΔELPD standard errors.

| Models | Rank ON | $\text{ELPD}_{\text{loo ON}}$ | $p_{\text{loo ON}}$ | $\Delta\text{ELPD ON}$ | $\text{SE}(\Delta\text{ELPD})$ ON | Weight ON | Rank OFF | $\text{ELPD}_{\text{loo OFF}}$ | $p_{\text{loo OFF}}$ | $\Delta\text{ELPD OFF}$ | $\text{SE}(\Delta\text{ELPD})$ OFF | Weight OFF |
| --- | --- | --- | --- | --- | --- | --- | --- | --- | --- | --- | --- | --- |
| PT GH | 1 | -522.05 | 56.00 | 0.00 | 12.77 | 0.547 | 2 | -508.54 | 55.09 | 0.92 | 12.75 | 0.423 |
| PT IH | 3 | -525.49 | 55.97 | 2.86 | 12.55 | 0.110 | 1 | -507.62 | 59.87 | 0.00 | 13.39 | 0.556 |
| PT | 2 | -523.57 | 43.57 | 1.51 | 12.51 | 0.379 | 3 | -517.88 | 45.03 | 10.26 | 13.48 | 0.015 |
| Null | 4 | -549.38 | 17.57 | 33.56 | 9.47 | 0.006 | 4 | -535.45 | 17.70 | 27.83 | 10.30 | 0.004 |
| Null PTL | 5 | 580.46 | 20.08 | 58.41 | 5.65 | $1.08 \cdot 10^{-14}$ | 5 | -587.47 | 16.35 | 79.85 | 5.15 | $1.04 \cdot 10^{-23}$ |

**Table 2.**
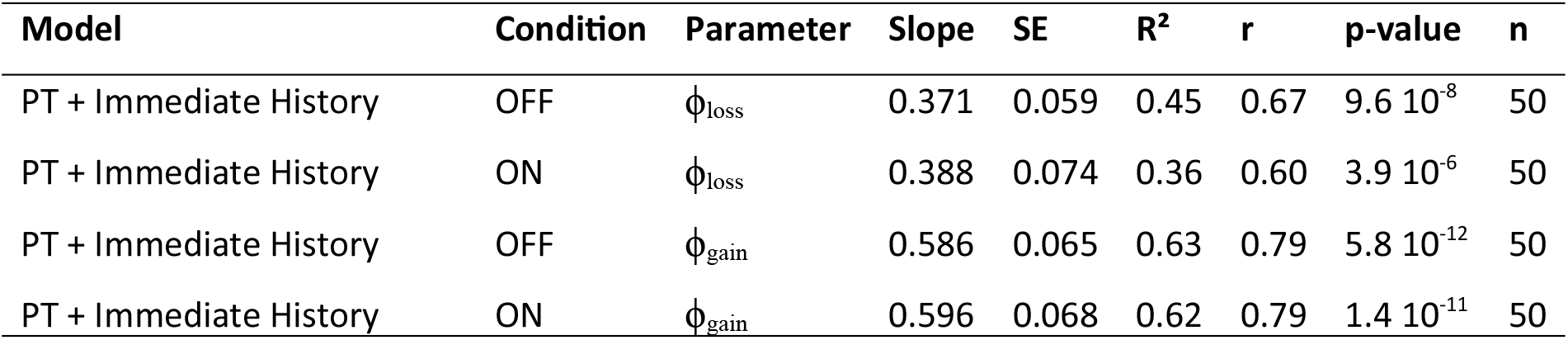
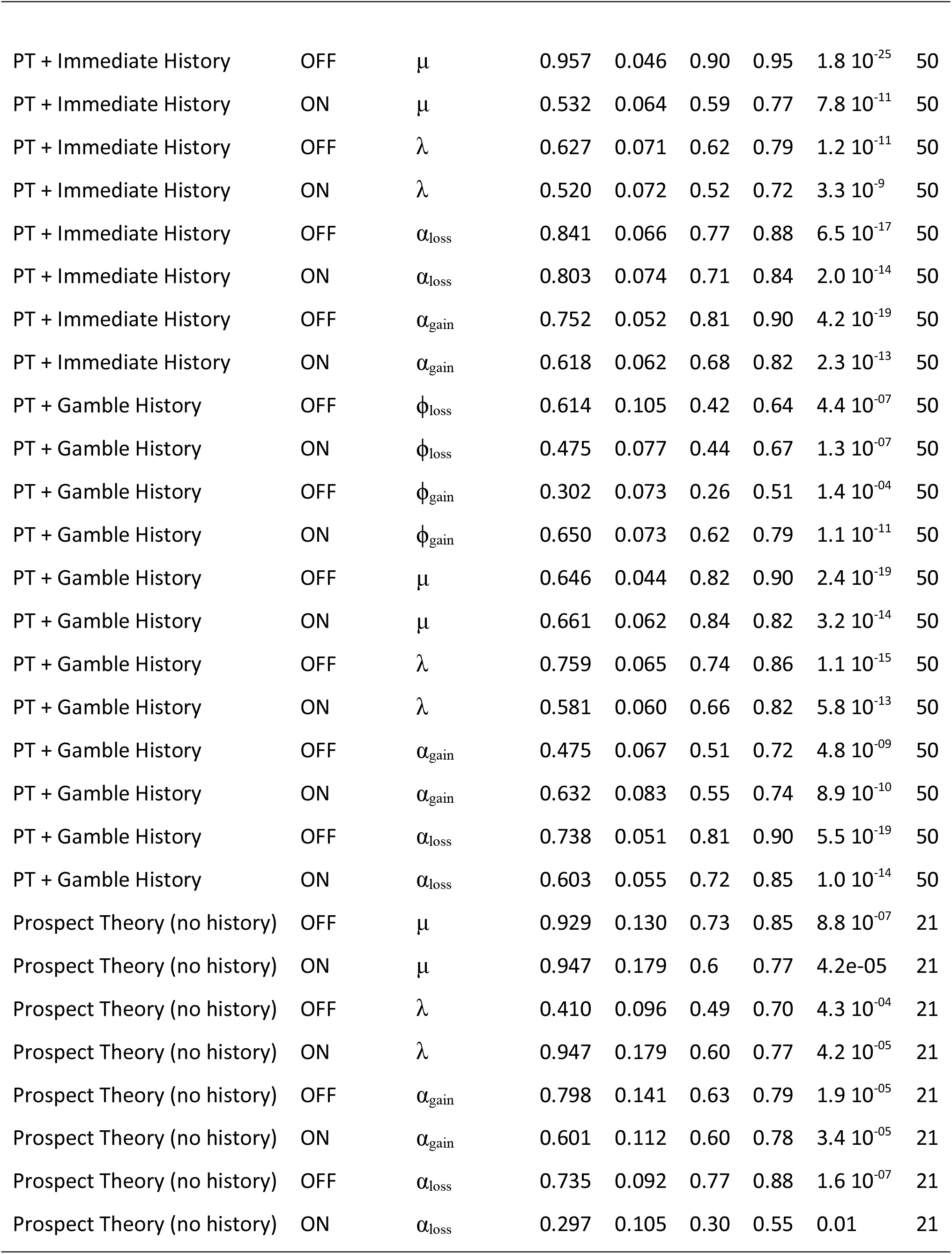
Parameter-recovery summary. Each row reports recovery accuracy for one parameter under one model and condition. “Model” names: Prospect Theory (no history) = PT; PT + Gamble History = PTL (uses the previous gamble’s outcome term); PT + Immediate History = PTL-Last (uses the immediately preceding trial’s outcome term). “Condition” indicates whether parameters used for simulation/fitting came from the ON or OFF stimulation estimates, **slope** ± SE and R^2^ from OLS (recovered ∼ true), p (two-tailed), and n (simulated units).

| Model | Condition | Parameter | Slope | SE | $R^2$ | $r$ | p-value | n |
| --- | --- | --- | --- | --- | --- | --- | --- | --- |
| PT + Immediate History | OFF | $\phi_{\text{loss}}$ | 0.371 | 0.059 | 0.45 | 0.67 | $9.6 \cdot 10^{-8}$ | 50 |
| PT + Immediate History | ON | $\phi_{\text{loss}}$ | 0.388 | 0.074 | 0.36 | 0.60 | $3.9 \cdot 10^{-6}$ | 50 |
| PT + Immediate History | OFF | $\phi_{\text{gain}}$ | 0.586 | 0.065 | 0.63 | 0.79 | $5.8 \cdot 10^{-12}$ | 50 |
| PT + Immediate History | ON | $\phi_{\text{gain}}$ | 0.596 | 0.068 | 0.62 | 0.79 | $1.4 \cdot 10^{-11}$ | 50 |
| PT + Immediate History | OFF | $\mu$ | 0.957 | 0.046 | 0.90 | 0.95 | $1.8 \cdot 10^{-25}$ | 50 |
| PT + Immediate History | ON | $\mu$ | 0.532 | 0.064 | 0.59 | 0.77 | $7.8 \cdot 10^{-11}$ | 50 |
| PT + Immediate History | OFF | $\lambda$ | 0.627 | 0.071 | 0.62 | 0.79 | $1.2 \cdot 10^{-11}$ | 50 |
| PT + Immediate History | ON | $\lambda$ | 0.520 | 0.072 | 0.52 | 0.72 | $3.3 \cdot 10^{-9}$ | 50 |
| PT + Immediate History | OFF | $\alpha_{\text{loss}}$ | 0.841 | 0.066 | 0.77 | 0.88 | $6.5 \cdot 10^{-17}$ | 50 |
| PT + Immediate History | ON | $\alpha_{\text{loss}}$ | 0.803 | 0.074 | 0.71 | 0.84 | $2.0 \cdot 10^{-14}$ | 50 |
| PT + Immediate History | OFF | $\alpha_{\text{gain}}$ | 0.752 | 0.052 | 0.81 | 0.90 | $4.2 \cdot 10^{-19}$ | 50 |
| PT + Immediate History | ON | $\alpha_{\text{gain}}$ | 0.618 | 0.062 | 0.68 | 0.82 | $2.3 \cdot 10^{-13}$ | 50 |
| PT + Gamble History | OFF | $\phi_{\text{loss}}$ | 0.614 | 0.105 | 0.42 | 0.64 | $4.4 \cdot 10^{-07}$ | 50 |
| PT + Gamble History | ON | $\phi_{\text{loss}}$ | 0.475 | 0.077 | 0.44 | 0.67 | $1.3 \cdot 10^{-07}$ | 50 |
| PT + Gamble History | OFF | $\phi_{\text{gain}}$ | 0.302 | 0.073 | 0.26 | 0.51 | $1.4 \cdot 10^{-04}$ | 50 |
| PT + Gamble History | ON | $\phi_{\text{gain}}$ | 0.650 | 0.073 | 0.62 | 0.79 | $1.1 \cdot 10^{-11}$ | 50 |
| PT + Gamble History | OFF | $\mu$ | 0.646 | 0.044 | 0.82 | 0.90 | $2.4 \cdot 10^{-19}$ | 50 |
| PT + Gamble History | ON | $\mu$ | 0.661 | 0.062 | 0.84 | 0.82 | $3.2 \cdot 10^{-14}$ | 50 |
| PT + Gamble History | OFF | $\lambda$ | 0.759 | 0.065 | 0.74 | 0.86 | $1.1 \cdot 10^{-15}$ | 50 |
| PT + Gamble History | ON | $\lambda$ | 0.581 | 0.060 | 0.66 | 0.82 | $5.8 \cdot 10^{-13}$ | 50 |
| PT + Gamble History | OFF | $\alpha_{\text{gain}}$ | 0.475 | 0.067 | 0.51 | 0.72 | $4.8 \cdot 10^{-09}$ | 50 |
| PT + Gamble History | ON | $\alpha_{\text{gain}}$ | 0.632 | 0.083 | 0.55 | 0.74 | $8.9 \cdot 10^{-10}$ | 50 |
| PT + Gamble History | OFF | $\alpha_{\text{loss}}$ | 0.738 | 0.051 | 0.81 | 0.90 | $5.5 \cdot 10^{-19}$ | 50 |
| PT + Gamble History | ON | $\alpha_{\text{loss}}$ | 0.603 | 0.055 | 0.72 | 0.85 | $1.0 \cdot 10^{-14}$ | 50 |
| Prospect Theory (no history) | OFF | $\mu$ | 0.929 | 0.130 | 0.73 | 0.85 | $8.8 \cdot 10^{-07}$ | 21 |
| Prospect Theory (no history) | ON | $\mu$ | 0.947 | 0.179 | 0.6 | 0.77 | $4.2 \cdot 10^{-05}$ | 21 |
| Prospect Theory (no history) | OFF | $\lambda$ | 0.410 | 0.096 | 0.49 | 0.70 | $4.3 \cdot 10^{-04}$ | 21 |
| Prospect Theory (no history) | ON | $\lambda$ | 0.947 | 0.179 | 0.60 | 0.77 | $4.2 \cdot 10^{-05}$ | 21 |
| Prospect Theory (no history) | OFF | $\alpha_{\text{gain}}$ | 0.798 | 0.141 | 0.63 | 0.79 | $1.9 \cdot 10^{-05}$ | 21 |
| Prospect Theory (no history) | ON | $\alpha_{\text{gain}}$ | 0.601 | 0.112 | 0.60 | 0.78 | $3.4 \cdot 10^{-05}$ | 21 |
| Prospect Theory (no history) | OFF | $\alpha_{\text{loss}}$ | 0.735 | 0.092 | 0.77 | 0.88 | $1.6 \cdot 10^{-07}$ | 21 |
| Prospect Theory (no history) | ON | $\alpha_{\text{loss}}$ | 0.297 | 0.105 | 0.30 | 0.55 | 0.01 | 21 |

## Supplementary Tables

**Supplementary Table 1.**
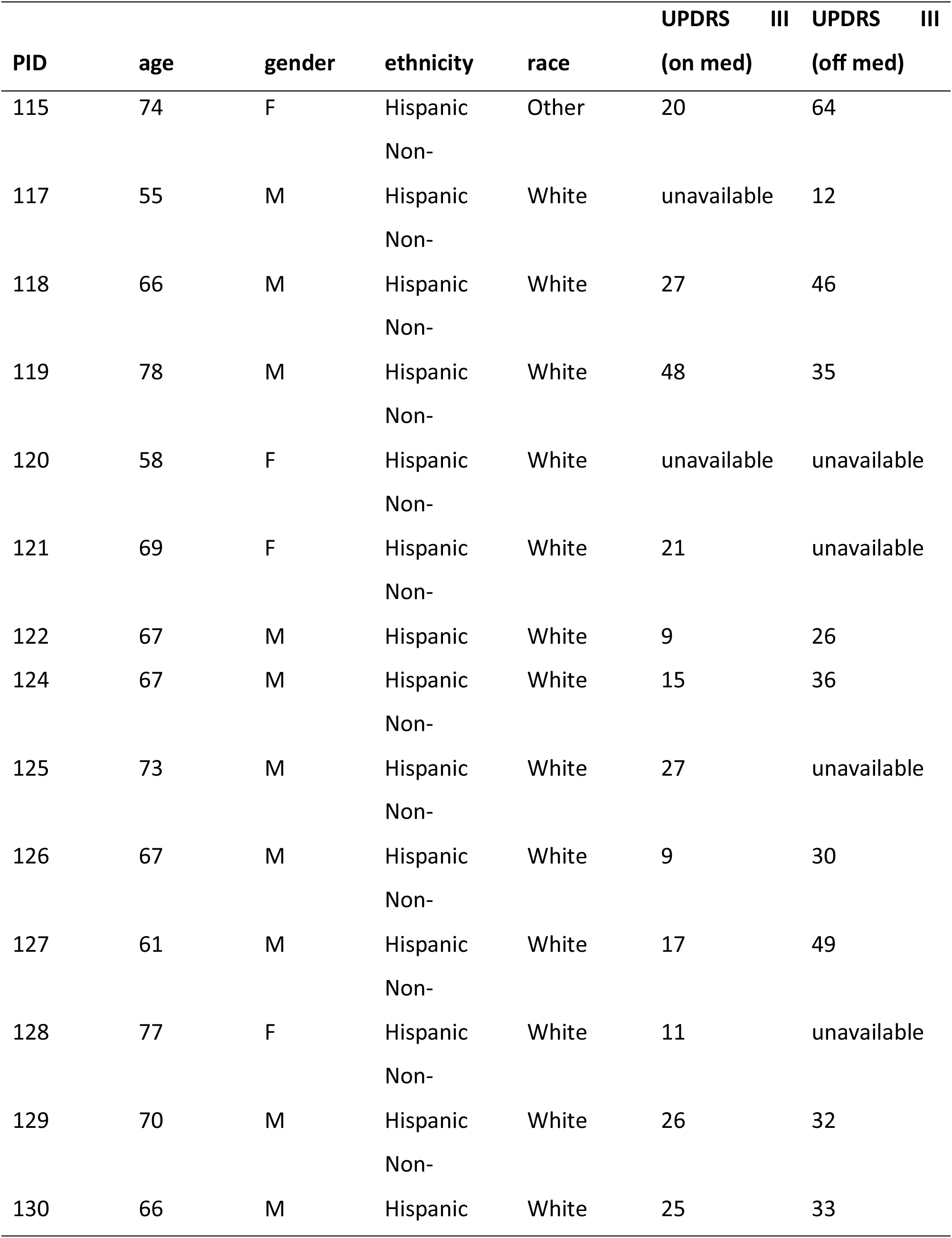
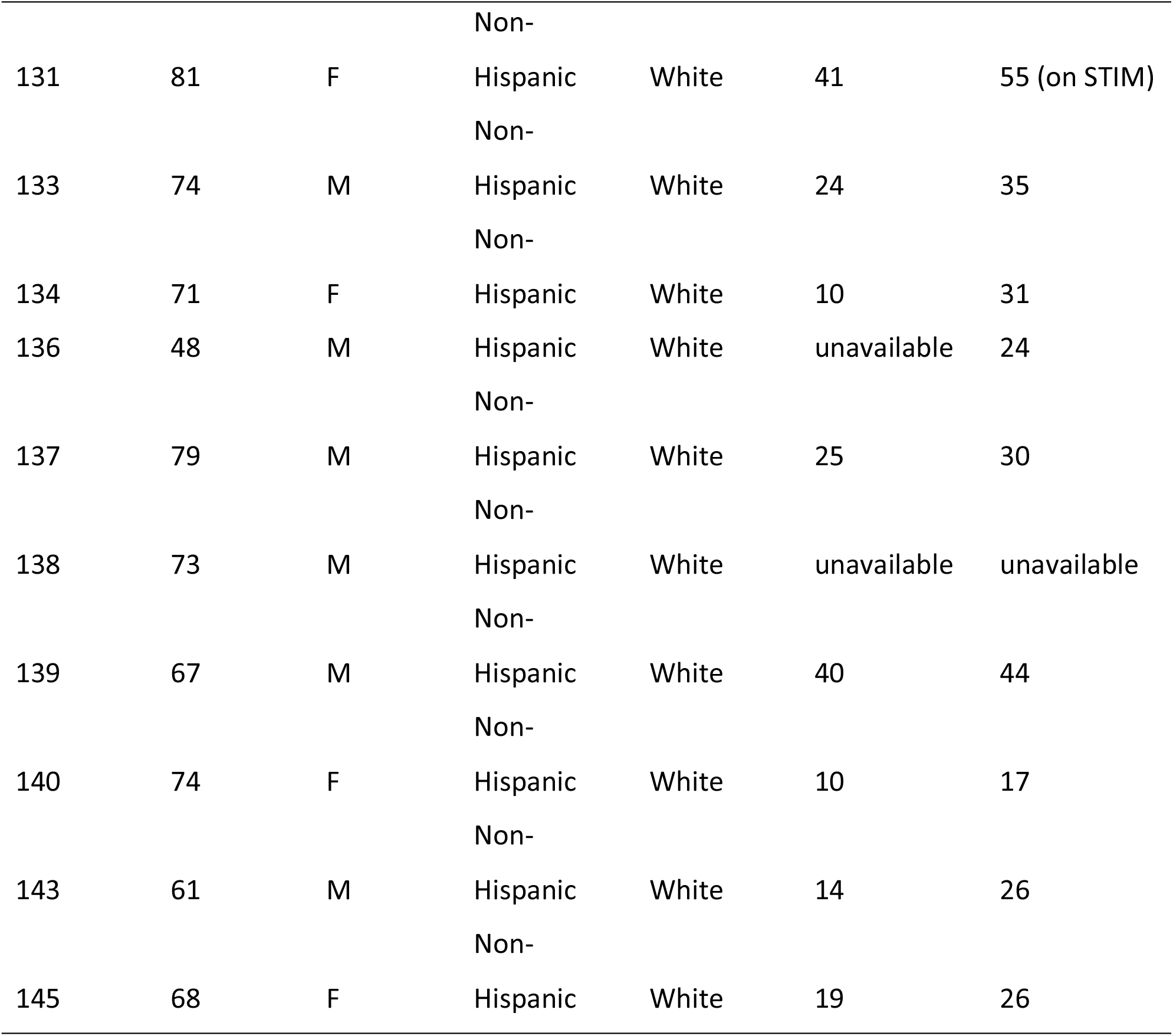
Patient Demographics and Clinical Characteristics. Summary of demographic and clinical information for each participant. The table includes patient ID, age at the time of surgery, gender, self-reported ethnicity and race, and Unified Parkinson’s Disease Rating Scale (UPDRS) Part III motor scores assessed before and after deep brain stimulation (DBS) surgery. These measures provide a clinical context for interpreting behavioral and neural results.

